# Synthetic control of actin polymerization and symmetry breaking in active protocells

**DOI:** 10.1101/2023.09.22.559060

**Authors:** Shiva Razavi, Felix Wong, Bedri Abubaker-Sharif, Hideaki T. Matsubayashi, Hideki Nakamura, Eduardo Sandoval, Douglas N. Robinson, Baoyu Chen, Jian Liu, Pablo A. Iglesias, Takanari Inoue

## Abstract

Non-linear biomolecular interactions on the membranes drive membrane remodeling that underlies fundamental biological processes including chemotaxis, cytokinesis, and endocytosis. The multitude of biomolecules, the redundancy in their interactions, and the importance of spatiotemporal context in membrane organization hampers understanding the physical principles governing membrane mechanics. A minimal, in vitro system that models the functional interactions between molecular signaling and membrane remodeling, while remaining faithful to cellular physiology and geometry is powerful yet remains unachieved. Here, inspired by the biophysical processes underpinning chemotaxis, we reconstituted externally-controlled actin polymerization inside giant unilamellar vesicles, guiding self-organization on the membrane. We show that applying undirected external chemical inputs to this system results in directed actin polymerization and membrane deformation that are uncorrelated with upstream biochemical cues, indicating symmetry breaking. A biophysical model of the dynamics and mechanics of both actin polymerization and membrane shape suggests that inhomogeneous distributions of actin generate membrane shape deformations in a non-linear fashion, a prediction consistent with experimental measurements and subsequent local perturbations. The active protocellular system demonstrates the interplay between actin dynamics and membrane shape in a symmetry breaking context that is relevant to chemotaxis and a suite of other biological processes.

## Introduction

Symmetry spans physics and biology. In physics, symmetry was first used to study crystalline structures.^1^ As the interplay between form and function has become more well-studied, applications of the concept of symmetry have broadened to other fields of physics and to living biological matter.^2,3^ In biology, processes such as chemotaxis, cell division, phagocytosis, and cell-cell fusion are driven by actin polymerization-induced forces that move the plasma membrane in single cells.^4–6^ Although the direction and magnitude of actin-generated forces can differ depending on the cellular processes at hand, symmetry breaking—a process in which a symmetric system exhibits directed behavior due to a bifurcation— has become a unifying hallmark of these processes.^7–10^ Asymmetric arrangement of actin, among other proteins, gives rise to spatial patterns that ultimately regulate cellular differentiation and development.^11–13^ Understanding how actin-induced asymmetries initiate and identifying their implications on function are foundational challenges that remain to be addressed in biophysics.^3,14,15^

In eukaryotic cells, multiple regulatory mechanisms, including protein switches and allosteric regulation of signal transduction, are known to amplify biomolecular asymmetries.^16, 17^ To study actin-induced symmetry breaking, in vitro reconstitution using reduced, membrane-bound systems that lack the biological complexity found in cells have provided insight into the components that regulate actin dynamics.^18–20^ Previous work has shown that cell-sized vesicles, placed in a mixture of actin and its polymerization regulators, experience F-actin-exerted forces on the membrane.^21–23^ These studies found that symmetry was broken in this system only if either capping protein (CP) or myosin were also present.^24–26^ Although they have been informative, these experiments have been limited to cases where actin and its regulators were external to the vesicles and in effectively infinite supply. Furthermore, these model systems were stochastic in how reactions proceeded, making it difficult to control these systems in space or time.^21, 22, 24, 25, 27, 28^

Here, we developed a synthetic platform for the programmable and chemically-inducible control of actin polymerization in giant unilamellar vesicles (GUVs). Using chemical protein dimerization modules as both sensing and actuation units, we show that rapamycin-induced recruitment of an engineered form of ActA, a potent Arp2/3-dependent actin nucleation promoting factor, to the inner leaflet of GUVs activates Arp2/3 complexes, leading to polymerization of G- and F-actin and generation of force on the membrane. We find that our platform couples biochemical cues to actin polymerization and exhibits chemically-induced symmetry breaking, wherein actin polymerization and GUV membrane deformations become asymmetric. Strikingly, when rapamycin is administered globally without a directional preference, the resulting asymmetric actin distributions and membrane deformations are uncorrelated with ActA distributions across GUVs, indicating symmetry breaking. Further microscopic analyses indicate that actin polymerization on the inner leaflets of GUVs results in membrane deformations that impart substantial shape eccentricity to GUVs. Modeling the dynamics of actin polymerization in GUVs, we find that the coupling of actin polymerization to asymmetrically-initiated actin nucleation sites is consistent with the empirically observed spread of actin. Thin-shell modeling further suggests that observations of GUV shape eccentricity are qualitatively consistent with the mechanical deformations imparted by an internal osmotic pressure, with the eccentricity determined by the degree of actin asymmetry. Together, these results demonstrate a synthetic biology platform and mechanistic model for the control of actin polymerization in GUVs, illustrating a tunable approach to the study of symmetry-breaking in self-organization and elucidating the interplay between biological signaling, actin dynamics, and thin-shell mechanics for shape generation.^29^

## Results

### Integrating sensing modules inside GUV protocells

To develop a synthetic biology-based platform that allows for control of actin polymerization and membrane remodeling in protocells in response to external chemical cues, we designed a protein dimerization-based sensing and actuation module inside GUVs (Supplementary Figure 1). We employed chemically-inducible dimerization (CID) based on FK506 binding proteins (FKBP) and FKBP-rapamycin binding domain (FRB) proteins, which heterodimerize in the presence of a small molecule derived from rapamycin. CID has been predominantly used in cells to manipulate biochemical reactions on the surface of membranes,^30^ tether artificial membranes,^31^ and mediate phase separation in emulsion,^32^ but to our knowledge has not been implemented inside GUVs.

We first individually linked FRB and FKBP to cyan and yellow fluorescent proteins, respectively (6xHis-CFP-FRB and 6xHis-YFP-FKBP). We encapsulated purified forms of these proteins inside GUVs composed of POPC and POPS membranes (see Table S1 for additional details) generated using the inverted emulsion technique.^33–35^ We found that stimulating the GUVs with an external, undirected supply of rapamycin resulted in heterodimerization of the FKBP and FRB proteins, as indicated by Förster resonance energy transfer (FRET) measurements of CFP. Rapamycin-induced heterodimerization completed within minutes; in contrast, administration of vehicle (dimethyl sulfoxide, DMSO) did not result in FRET signal intensity increase, indicating no heterodimerization (Supplementary Figure 1). Our observations verified that our engineered CID modules enable externally-driven signal processing inside GUVs.

To allow for membrane-localized output, we next designed CID-based constructs targeted to GUV membranes. We aimed to concentrate the luminal CFP-FRB at the inner membrane where FKBP was bound in a rapamycin-inducible manner (**Fig. 1A**). To this end, we coupled FKBP to the effector domains of the myristoylated alanine-rich kinase substrate (MARCKS-ED), a positively-charged peptide that binds to phosphatidylserine (PS).^36^ The resulting construct, mCherry-FKBP-MARCKS, and luminal CFP-FRB were purified, introduced into PS-containing GUVs (Table S1), and confirmed to be present at designated GUV locations (**Fig. 1B**). Upon rapamycin administration, the luminal CFP-FRB localized to membranes (**Fig. 1B,C** and movie S1) within minutes, a timescale similar to that previously observed with CID in mammalian cells.^37^ The minute-scale temporal dynamics of our GUV-encapsulated CID system suggests that it can recapitulate actin’s temporal dynamics in natural cellular environments.

**Figure 1.**
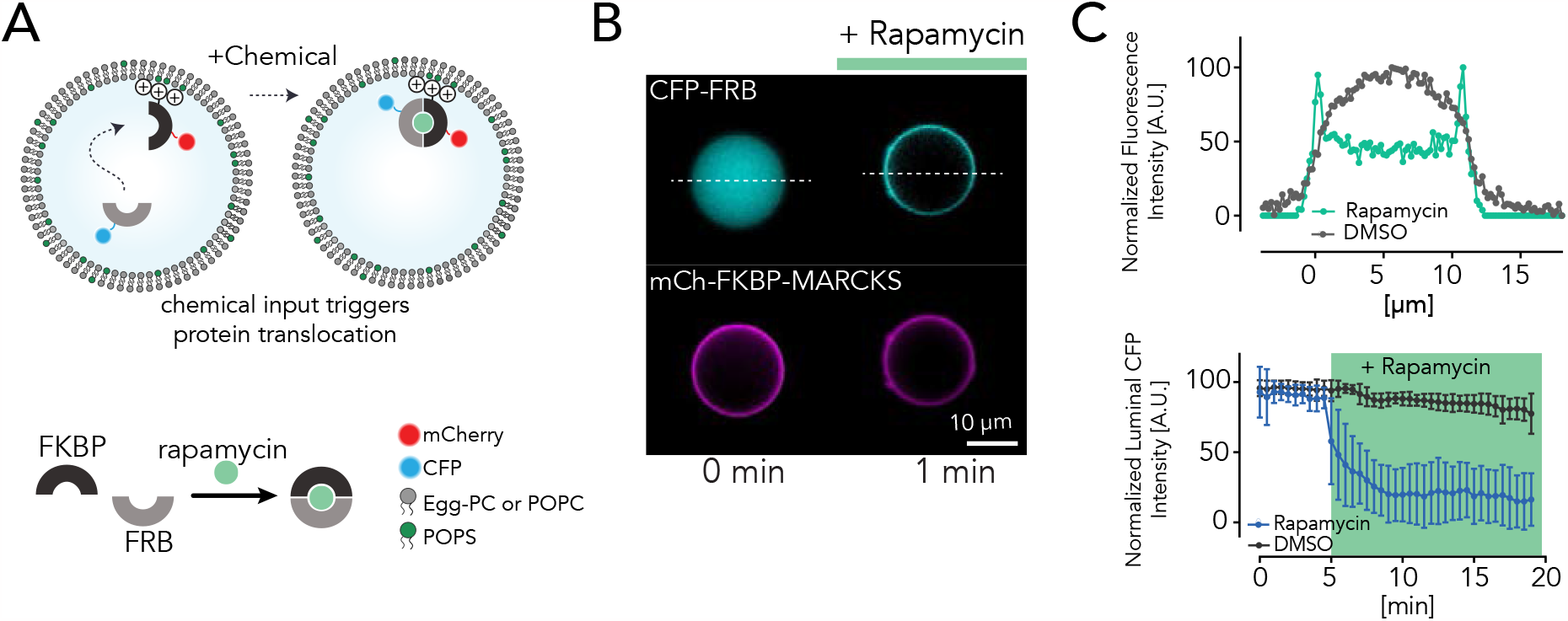
Reconstitution of actuation inside protocells. **A**. Schematic of the GUVs containing the membrane-anchored mCh-FKBP-MARKCS and luminal CFP-FRB proteins. Rapamycin-induced dimerization of FKBP and FRB moves the CFP-FRB protein towards the membrane. **B**. Epi-fluorescence images of CFP-FRB (4.4 *μM*) and mCh-FKBP-MARKCS (7.8 *μM*) in the symmetric GUVs containing POPC: POPS (4:1 mol%) in the inner lipid leaflet. The initially luminal CFP-FRB protein switches localization to the membrane within minutes post-rapamycin administration.**C**. Line scans of the mCherry and CFP fluorescence intensity across the vesicle show protein localization before and after rapamycin addition (top). The luminal fluorescence intensity of the CFP-FRB (bottom) normalized by the average of the initial values prior to rapamycin treatment is plotted, highlighting the minute-scale CFP-FRB translocation towards the membrane only in the presence of rapamycin but not DMSO vehicle. n=14 for both rapamycin and DMSO conditions. Error bars represent standard deviation. The green box marks rapamycin presence.

### Inducible, on-demand actin polymerization inside GUVs

We next coupled our CID-based constructs targeted to GUV membranes with actin polymerization. We linked FRB to a re-engineered domain of actin assembly-inducing protein, ActA (1-183). ActA is a potent Arp2/3-dependent actin nucleation promoting factor^38, 39^ that we recently engineered to generate forces on the cytoplasmic membranes of mammalian cells.^40^ We encapsulated a purified, C-terminal 2xstrep affinity-tagged form of the ActA-FRB construct, 2xstrep-ActA-FRB-CFP, together with purified Arp2/3 complex, purified G-actin mixed with 8% Alexa Fluor 488-labeled G-actin for visualization, purified 6xHis-tagged mCherry-FKBP, and ATP in GUVs containing nickel-conjugated lipids to enable His-tagged FKBP anchoring (**Fig. 2A**, Table 1, Supplementary Figure 2, and Table S1).

**Table 1.** Biomolecular modules used in the active protocells.

|  | Part | Function | Concentration |
| --- | --- | --- | --- |
| Protein Network | 6xHis-mCherry-FKBP | chemical sensing - membrane anchoring | 9.5 μM |
| | ActA-FRB-CFP | chemical sensing - Arp2/3 activation | 2-8.5 μM |
| | Arp2/3 complex | actin nucleation | 0.15 - 0.5 μM |
| | G-actin | force generation | 1.5-3 μM |
| Lipids | POPC | biological membrane mimic | 94 mol% |
|  | DGS-NTA(Ni) | anchoring sensory modules | 4 mol% |
|  | PEG | reducing unspecific lipid-protein interaction | 1 mol% |
| Inner Buffer | ATP | energy source | 1 mM |
|  | MgCl<sub>2</sub> | actin stability and polymerization | <1 mM |
|  | KCl | physiological salt | <50 mM |
|  | Tris-HCl (pH 7.5) | pH stability | >10 mM |
|  | sucrose | osmotic pressure balance | 850 mM |

**Figure 2.**
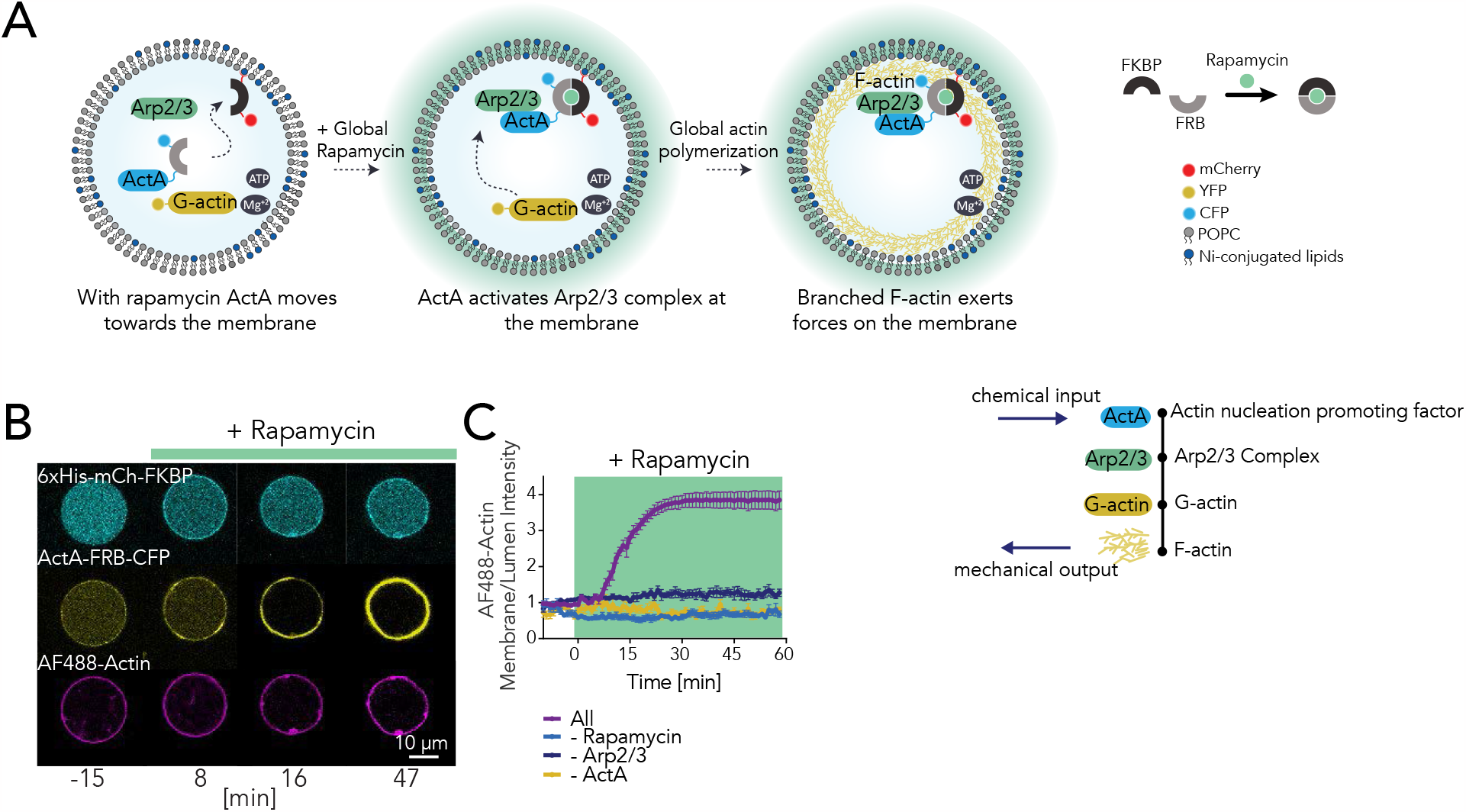
Reconstitution of actin-induced force generation inside rapamycin sensing active protocell. **A**. Schematic of the signaling pathway. ActA activates Arp2/3 complex on the membrane, where F-actin branches grow to generate force. CID system modules coupled with ActA (1-183) transduce rapamycin sensing to force actuation. ActA (1-183)-FRB-CFP, Arp2/3, and G-actin are initially diffuse in the lumen. mCherry-FKBP is anchored at the membrane.**B**. Confocal images of symmetric POPC: DGS-NTA(Ni): PEG-PE (94:4:1) GUVs containing 6xHis-mCh-FKBP, ActA (1-183)-FRB-CFP (2 *μM*), Arp2/3 (150 *nM*), G-actin (3 *μM*), and ATP (1 *mM*) in Mg^2+^ buffer. All protein constructs except 6xHis-mCh-FKBP are luminal at t=0. With rapamycin (10 *μM* final concentration), ActA translocates to the membrane and triggers F-actin forces. **C**. Time-course of mean actin localization in the presence of rapamycin and all components of the signaling cascade, as well as negative controls where Arp2/3, ActA, or rapamycin are absent. n=6 for All, n=4 for -Arp2/3, n=9 for -ActA, and n=10 for DMSO conditions. Error bars indicate standard error of the mean(SEM). Green area marks rapamycin presence.

With this design, initially all components, except for membrane-bound mCherry-FKBP, were inside the luminal body. After administration of rapamycin to the external milieu, we found that the engineered ActA redistributed itself on the membrane and activated the Arp2/3 complex, as supported by actin assembly measurements in test tubes (Supplementary Figure 3). Subsequently, stochastic and asymmetric patches of actin emerged on the membrane and grew to engulf the GUV boundary (**Fig. 2B** and movie S2). With sufficiently high concentrations of G-actin (*∼*3 *μM*), actin polymerization on the membrane deformed GUVs smaller than *∼*50 microns in diameter (**Fig. 2B**). To validate that membrane deformations emerge as a result of actin assembly, we externally administered latrunculin A, a toxin that depolymerizes F-actin, binds G-actin, and hinders actin assembly.^41^ With latrunculin A present in the media, we found that actin patches failed to grow; additionally, the fluorescence intensities of already-developed F-actin patches decreased over time (Supplementary Figure 4). These results indicate that actin growth was caused by actin polymerization, rather than the accumulation of monomeric G-actin. Indeed, without ActA or Arp2/3, actin failed to polymerize. In the absence of rapamycin, we found that only a baseline level of background luminal actin polymerization occurred (**Fig. 2C** and Supplementary Figure 5). These observations suggest that a threshold concentration of ActA drives actin polymerization, a finding similar to previous results indicating that the WASP family of Arp2/3 activators^42^ also exhibits a concentration threshold for downstream Arp2/3 activation. Notably, our rapamycin-inducible control of a threshold-dependent signaling cascade, which leads to actin polymerization, provides a system that can explore biological symmetry breaking using only a small number of regulators.

### Actin polymerization is dynamic and exhibits symmetry breaking

In order to quantify the dynamics of actin polymerization in our model system, we generated kymographs for the spatial distributions of ActA and actin fluorescence intensities, as well as membrane curvature, as functions of time (methods and supplementary materials) (**Fig. 3A**). These kymographs showed that, in typical GUVs, ActA and actin were homogeneously distributed in space until a timescale of *∼*5 min after rapamycin administration (at *t* = 0), after which discrete actin nucleation sites formed over the membrane surfaces. Principal component analyses further showed that multiple discrete peaks in the actin and local curvature kymographs, and to a substantially lesser extent in the ActA kymographs, accounted for most of the empirically observed variation across GUVs (Supplementary Figure 6). To investigate whether the heterogeneity of actin nucleation was associated with upstream ActA distribution, we calculated ActA-actin correlations for each of 18 GUVs across time (**Fig. 3B**). These analyses indicated a small ActA-actin correlation of *∼*0.2, which likely arose from lipid artifacts in the membrane where proteins appeared enriched. Importantly, these correlations remained unchanged after the addition of rapamycin. However, discrete and asymmetric actin nucleation occurred after rapamycin addition, as supported by plots of actin fluorescence intensity along the GUV contours (**Fig. 3C**). The absence of correlation increase with respect to ActA therefore indicates that the symmetry breaking is spontaneous and does not depend on detectable levels of ActA, a finding consistent with observations of symmetry breaking in previous studies.^43^

**Figure 3.**
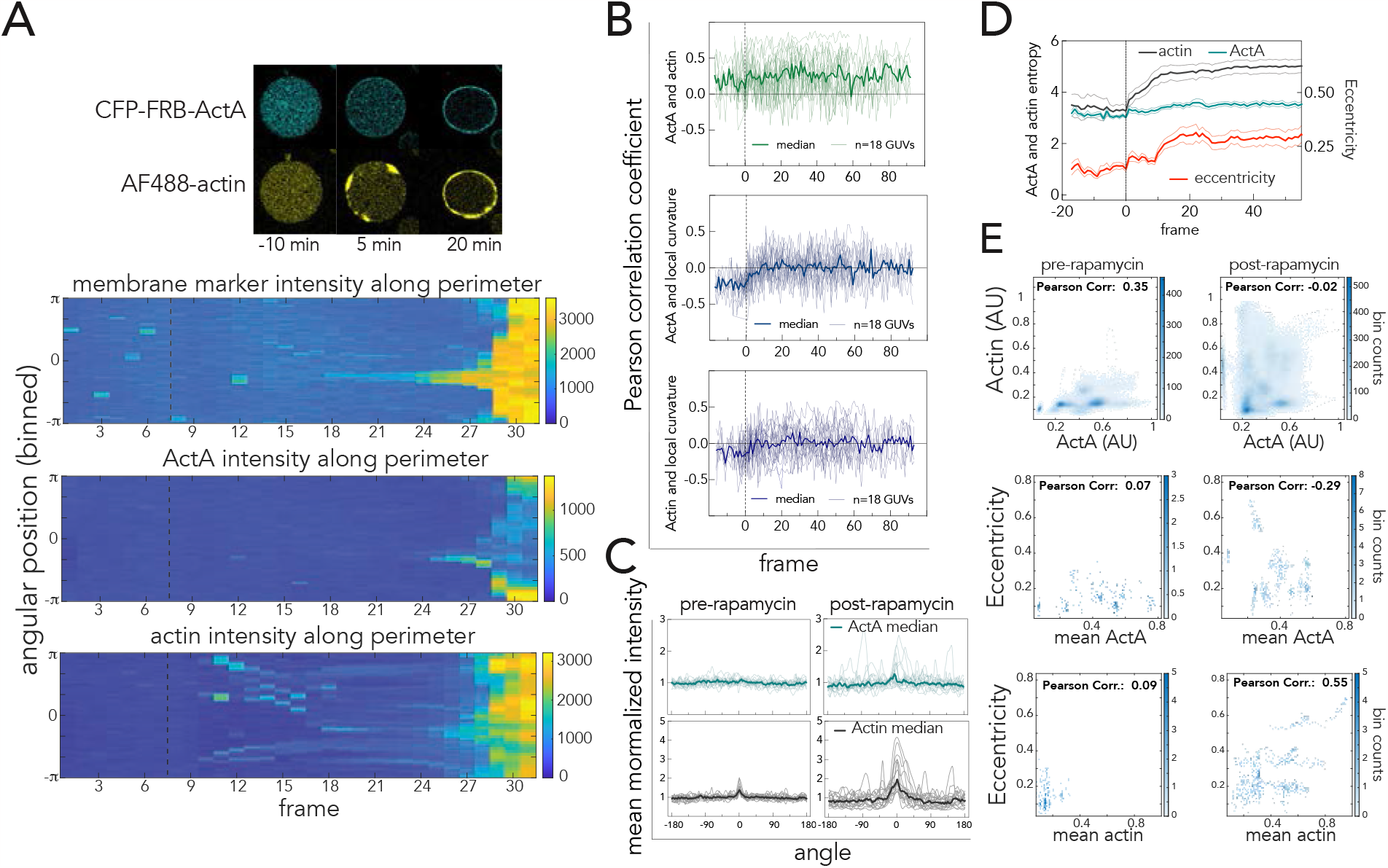
Symmetry breaking in biochemical and physical spaces on a local and global scale. **A**. Representative confocal image of a GUV before and after rapamycin addition together with its corresponding kymographs tracking enrichment of ActA and actin on the membrane in time. t=0 and the dashed black lines on the kymographs represent the rapamycin addition timepoint. **B**. Tracking the Pearson correlation coefficient between biochemical parameters (ActA and actin) over time highlights no correlation change upon rapamycin addition (top). The Pearson correlation coefficient evolution for ActA and local curvature become further decorrelated upon rapamycin addition (middle). Similarly for actin and local curvature, no correlation is observed (bottom). **C**. Mean-normalized fluorescence intensity profiles of ActA and actin shown at two frames pre-rapamycin and 10 frames post-rapamycin addition. **D**. Plots of the entropy of boundary parameters over time, highlighting that ActA entropy remains steady while those of actin and eccentricity both spike after rapamycin addition. Error bars indicate SEM. **E**. Plots of mean ActA intensity, mean actin intensity, and eccentricity, where the bulk correlation is computed over aggregated data from all GUVs for 18 frames pre-rapamycin and 35 frame post-rapamaycin. For each plot, n= 18.

We next performed similar correlation analyses for membrane curvature (**Fig. 3B**). These analyses indicated that ActA and actin were essentially uncorrelated with membrane curvature, despite a small increase in correlation from −0.2 to 0 after rapamycin addition for ActA and curvature (**Fig. 3B**). While actin polymerization may lead to local membrane deformations, it is possible that these deformations may manifest in membrane curvature alterations on a global scale. To investigate this hypothesis further, we quantified the entropy as a global measure of variation—reflecting changes in the distributions of values, such that higher entropy values indicate higher asymmetry—across all points on the GUV contours (**Figs. 3C,D**). We calculated the entropy for ActA intensity and actin intensity; additionally, we calculated the eccentricity of the best-fitting ellipse to each GUV contour as a global measure of shape eccentricity. We found that, in contrast to ActA entropy, the actin entropy and eccentricity persistently increased after rapamycin addition. These results are consistent with our observations that actin nucleation is asymmetric and uncorrelated with ActA (**Figs. 3B,D**), and further suggest that asymmetry in the actin distribution is associated with global, but not local, GUV shape changes.

While we have focused on the correlations between ActA, actin, and curvature for each GUV, these correlations may differ globally across GUVs. To investigate this possibility, we computed ActA and actin mean intensity profiles at representative timepoints pre- and post-rapamycin treatment (**Fig. 3E**). Intriguingly, and similar to our previous analyses (**Fig. 3B**), we found no correlation between the mean intensities of ActA and actin after rapamycin treatment, despite the two quantities being moderately positively correlated—possibly due to initial lipid artifacts in the membranes—before rapamycin treatment. Correlations between mean ActA intensity and eccentricity were weaker, while mean actin intensity was uncorrelated with eccentricity before rapamycin treatment. The strongest correlation, of *∼*0.55, emerged between mean actin intensity and eccentricity after rapamycin treatment, again indicating that increased actin polymerization is associated with asymmetric GUV shapes. Taken together, these results further highlight symmetry breaking in actin polymerization and GUV shape in our synthetic platform.

### Actin polymerization is associated with GUV shrinkage and shape eccentricity

As our analyses indicate that asymmetric actin polymerization is associated with changes in GUV shape, we further investigated the dynamical and mechanical implications of actin symmetry breaking. Building on our observations of actin fluorescence at the GUV surface across all timepoints, we assumed that a spherical, linear-elastic, and isotropic thin-shell layer of actin is initially anchored to the inner leaflet of each GUV, and that the membrane surface area is larger than that of the actin shell’s. As internal, outward turgor pressure on the order of *∼*0.1 to 1 atm can be generated by the millimolar-scale osmotic imbalance of solutes across the GUV (Table 1), this actin shell is the main load-bearing element, and will be remodeled by additional actin polymerization (**Fig. 4A**).

**Figure 4.**
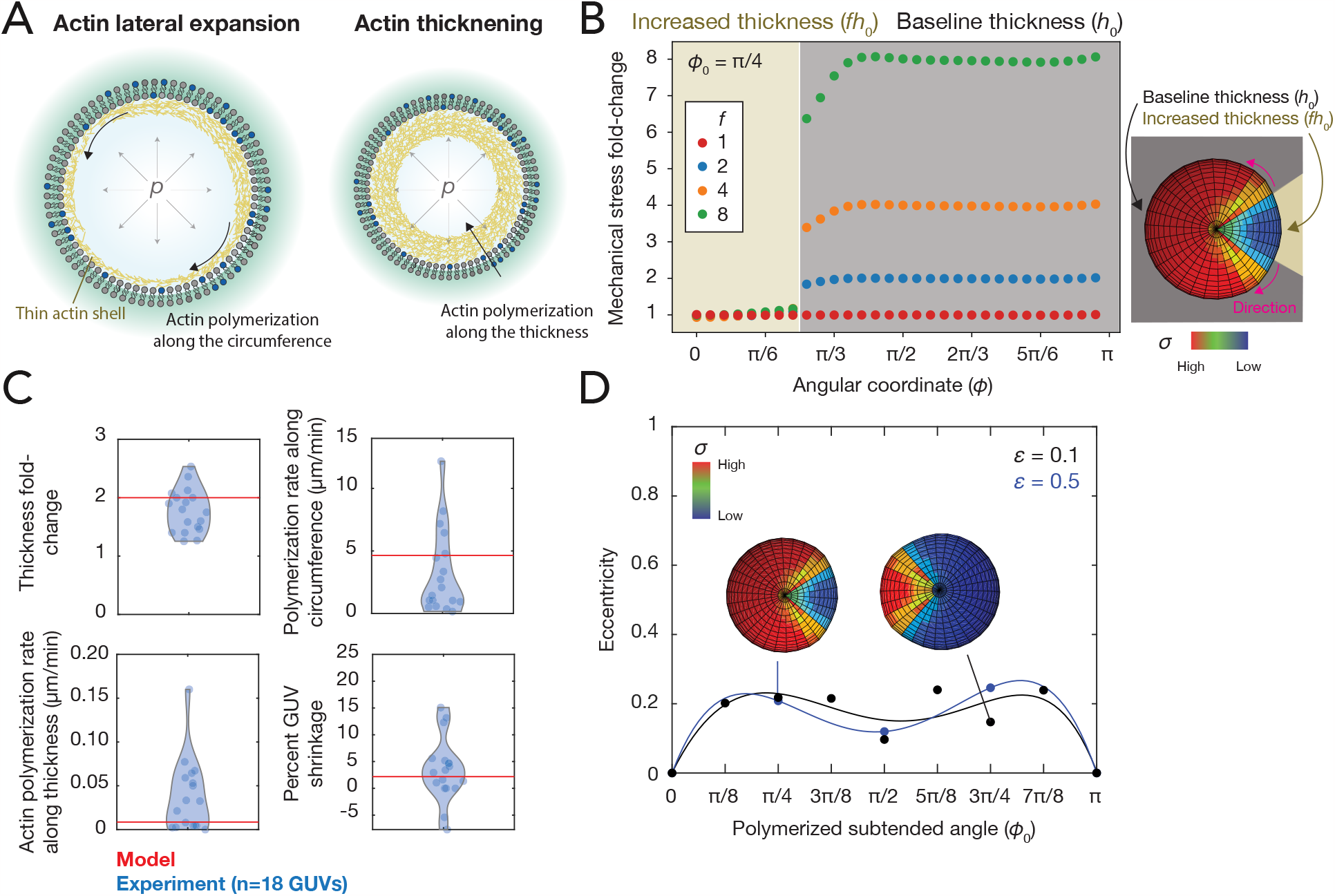
Biophysical modeling of the dynamics and mechanics of actin polymerization. **A**. Schematic of actin dynamics predicted by the model, in which the volumetric rate of actin polymerization depends on the mechanical stress. Actin polymerization is predicted to predominantly occur along the circumference (lateral expansion), then along the GUV thickness (thickening). The actin is viewed as a thin elastic shell loaded by the internal turgor pressure, *p*. **B**. Plot of the fold-change in mechanical stress, *σ*(*φ*)/*σ*_0_, where *σ*(*φ*) is the average maximum principal stress at the angular coordinate *φ* and *σ*_0_ = *pr*_0_/2*h*_0_ is the principal stress where the thickness is *h*_0_. Results are shown for a polymerized subtended angle of *φ*_0_ = *π*/4 and different thickness factors of *f* = 1, 2, 4 and 8. (Inset) Representative finite-element simulation results. The maximum principal stress is visualized. **C**. Comparison of model predictions (red lines) with experimental measurements (blue points) for the 18 GUVs analyzed in Fig. 3, for characteristic parameter values of *h*_0_ = 0.1 *μm, f* = 2, = 0.1 *μm, r*_0_ = 10 *μm, v*_0_ = 200 min^*−*1^, actin elastic modulus *E* = 0.1 GPa,^57, 58^ actin Poisson’s ratio *v* = 0, and turgor pressure *p* = 1 atm. Rates were calculated as linear approximations; see Supplementary Information for details. **D**. Plot of simulated GUV eccentricity values as a function of the polymerized subtended angle, *φ*_0_, for the same parameter values as in **C**, corresponding to mechanical strains of *pr*_0_/*Eh*_0_ = 0.1 (black). A value of *p* = 5 atm was also used in the simulations, corresponding to mechanical strains of 0.5 (blue).

Previous studies have assumed that the actin polymerization rate outside a droplet, with actin polymerization occurring in one direction, depends on the normal stress in the droplet with a Kramer’s rate dependence.^44^ Here, building on this assumption of mechanical stress-dependent actin polymerization, we assume that the local, volumetric rate of actin polymerization (*dV* /*dt*) at any surface coordinate (*θ, φ*) and at time *t* depends on the in-plane mechanical stress, *σ*(*θ, φ, t*), also as a Kramer’s rate: 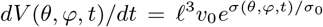, where *σ*_0_ is a baseline value of stress, . *ℓ* is the length of an actin monomer, and *v*_0_ is a unit rate of actin polymerization. For a spherical actin shell, the nonvanishing in-plane stresses are *σ*_0_ = *pr*_0_/2*h*_0_, where *p* is the turgor pressure, *r*_0_ is the shell radius, and *h*_0_ is the initial actin shell thickness. Anticipating that ActA-dependent actin polymerization results in the local shell thickness being multipled by a factor of *f≫*1, we expect that the ActA-enriched regions with highest mechanical stress are at the leading edges; this is confirmed by finite-element simulations of pressurized spherical shells with two different thicknesses (**Fig. 4B** and Supplementary Information). When actin finishes polymerizing laterally across the GUV surface at time *t*^*i*^, the stresses are approximately homogeneous and the form of *dV* /*dt* suggests that actin thickens linearly in time. Detailed calculations for this model (Supplementary Information) predict that:

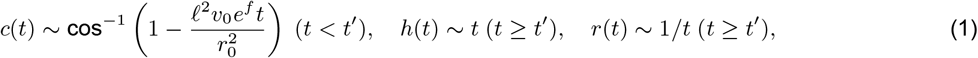

where *c*(*t*) is the fraction of the shell circumference at which ActA-dependent actin polymerization has already occurred at time *t, h* is the polymerized thickness of the actin shell at time *t*, and *r* is the radius of the actin shell at time *t*, which decreases due to shell thickening counteracting the turgor pressure and decreasing the mechanical strain, *ε*, in the actin shell (supplementary information). For characteristic values of all other parameters, fitting the unit actin polymerization rate (*v*_0_) predicts both the quantitative rates of actin’s lateral expansion and thickening, and we find that these rates and the resulting shrinkage of the actin shell are consistent with our empirical observations (**Fig. 4C**). Furthermore, quantifying the magnitude of GUV shrinkage toward the end of our timelapses reveals a radial shrinkage on the order of *∼* 5%, consistent with our model predictions (**Fig. 4C**).

The consistency between our empirical observations and our biophysical model therefore suggests that asymmetric actin polymerization is a dynamical process that results in mechanical alterations to GUVs. Moreover, at long timescales, actin thickening can result in GUV actuation. Given that actin symmetry breaking was also associated with shape eccentricity (**Fig. 3D**), we further investigated whether mechanical deformations could give rise to substantial shape eccentricity. For simplicity, we modeled actin as a shell with two thicknesses, using finite-element simulations across a range of angles subtended by the thicker actin (**Fig. 4D**). We found that, for finite values of mechanical strain and across a broad range of shell thicknesses, the shape of the simulated actin shell was eccentric and could be fit by ellipsoids with eccentricity values on the order of *∼*0.2 (**Fig. 4D**), consistent with the magnitude of eccentricity values inferred from experiments (**Fig. 3D**). These results indicate that shape asymmetry can arise as a mechanical consequence of actin symmetry breaking alone.

### Asymmetric inputs validate time-scales and elucidate the origins of symmetry breaking

In order to further probe our system, we reduced its spatial degrees of freedom by constraining the position of the rapamycin input. We induced a rapamycin gradient on the length-scale of a GUV using a point source by administering ethanol-solubilized rapamycin with a microinjector close to each GUV and adding Alexa Fluor 647 dye to track the rapamycin gradient (as its Stokes radius is comparable to that of rapamycin^45^). In order to confine the localization further, we used deformable DPPC lipids, which exhibit lower lateral diffusion and membrane fluidity as compared to POPC lipids and still deform upon actin polymerization^46^ (Supplementary Figures 7,8 and table S1). Additionally, with all other conditions held constant, we reduced the ActA concentration to limit its availability (**Fig. 5A** and Supplementary Table S2). After administering rapamycin locally, we observed that ActA always translocated towards the membrane region proximal to the needle tip, and within *∼*5 minutes, ActA decorated the entire GUV membrane. Furthermore, we found that subtle, but stable, ActA localization at the needle tip, on the order of *∼*1 minute, was sufficient to bias an actin nucleation site to appear at the same site. These observations are consistent with the actin nucleation dynamics observed in an in vitro pyrene assay (Supplementary Figure 3). Intriguingly, transient ActA leading to stable actin accumulation is reminiscent of memory behavior due to time delay,^47^ and these observations indicate that ActA may store positional information downstream of the local amplification potentially mediated by the Arp2/3 complex (Supplementary Figure 9).

**Figure 5.**
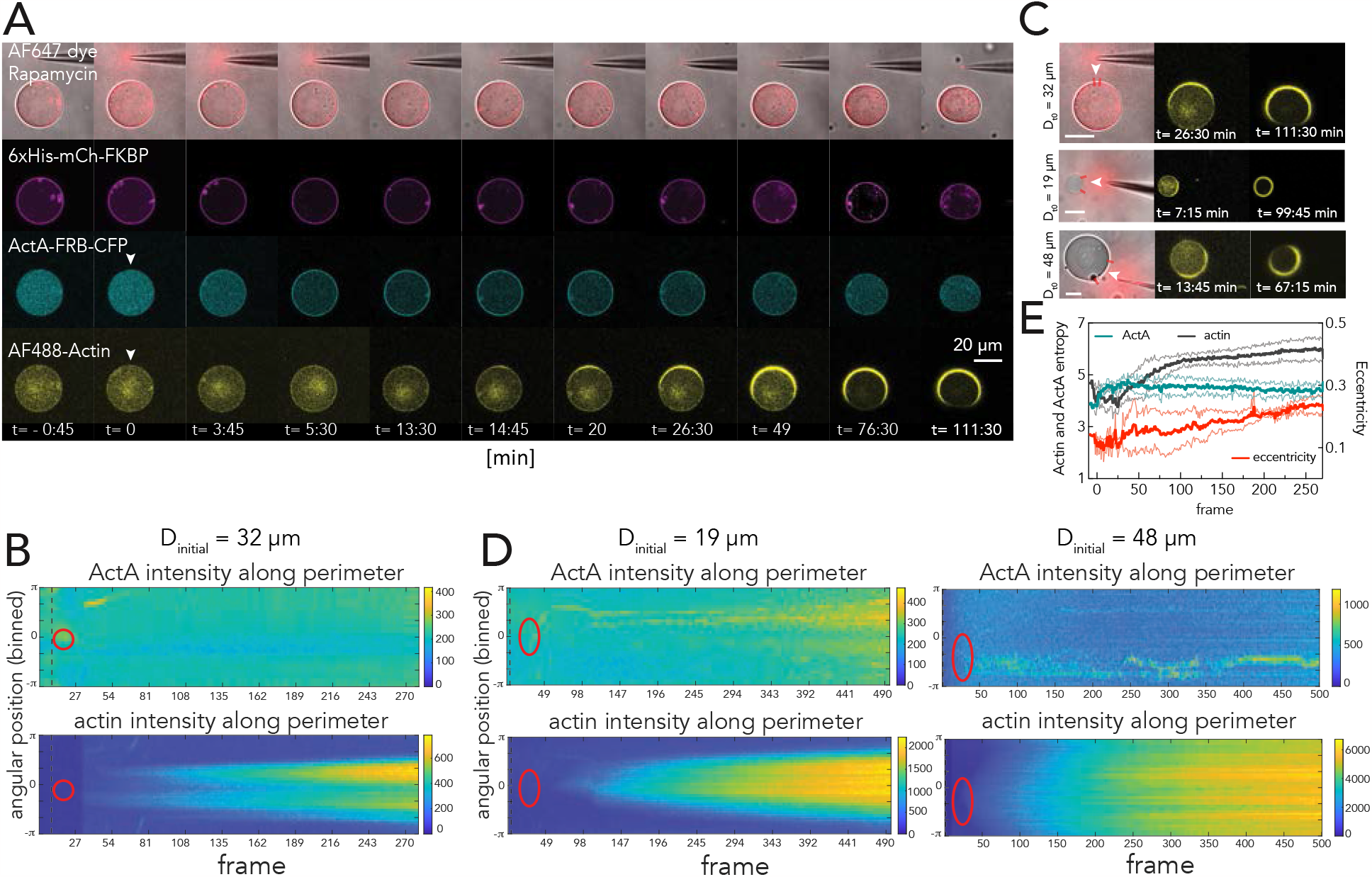
Reconstitution of spatially controlled localized actin deformation. **A**. Confocal images of GUVs during rapamycin administration. Upon rapamycin release (marked by the pink Alexa fluor 647 dye signal), ActA first translocates to the location marked by the white arrow. Local F-actin appears at the same site within *∼*15 min. **B**. Kymographs showing the evolution of the ActA and actin signal intensity on the membrane for the GUV presented in (A). The red oval marks the effective high rapamycin concentration area. **C**. Comparison of the actin thickening versus spreading, for GUVs of different radii. The area between the two lines highlights the effective rapamycin area. Scale bar is 20 *μm*. **D**. Similar to (B), but showing a kymograph of the signal accumulation for the 48 *μm* and 19 *μm* GUVs depicted in (C). Time 0 corresponds to the rapamycin input addition. **E**. Plots of the entropy of boundary parameters over time, highlighting that ActA entropy remains steady while that of actin and eccentricity both spike after rapamycin addition. Error bars indicate SEM.

Downstream of transient ActA localization, we found that, within *∼*15 minutes—by which time the ActA distribution appeared uniform across the membrane—F-actin patches emerged next to the needle tip and grew to encompass *∼*85% of the membrane (**Fig. 5B,C**, and movie S3), expanding laterally and in thickness similarly to our previous experiments, in which rapamycin was administered without a directional bias. These results show that asymmetric actin polymerization is uncorrelated with ActA irrespective of the route (undirected or directed) of rapamycin induction. As in the previous experiments, substantial shape eccentricity and membrane shrinkage also occurred, consistent with our model predictions (**Figs. 5E, 4D**). Additional perturbations support that actin symmetry breaking is associated with local fluctuations in rapamycin concentration, and that actin polymerization is indeed required for membrane shape deformations: when we altered the needle placement in independent experiments, we observed a positive correlation between the needle tip and actin polymerization initiation sites. This manipulation ruled out the possibility of flow-based interference or phase-separated lipids driving the local output (**Figs. 5C,D**), and further suggested that local, non-equilibrium fluctuations in rapamycin concentration—which may result in small ActA fluctuations not seen in our correlation analyses—is associated with actin symmetry breaking **Fig. 5E**. Consistent with this hypothesis, we found enhanced localization of F-actin after increasing the GUV diameter size, suggesting that symmetry breaking is length-scale dependent (**Figs. 5B,C,D** and Supplementary Figure 10). For larger GUVs, the relatively large membrane surface adjacent to the needle likely depletes the reaction substrate and serves as an internal non-molecular inhibitory element.

We last investigated the factors influencing actin thickening and GUV shape deformations (**Fig. 5D**). To assess if we can sculpt this thickening by integrating additional actin modulators, we used a purified capping protein, Cap32/34, which binds the barbed end of actin, preventing actin monomer addition or loss. In the presence of 100 *nM* of purified Cap32/34,^48^ we found that actin patches became sparse and thin and eventually disintegrated. This indicated that actin polymerization at barbed ends underlied actin thickening, consistent with bulk actin polymerization assay measurements (Supplementary Figure 3) in which actin polymerization was inhibited with overloaded Cap32/34 (3 *μM* ; Supplementary Figure 11). Moreover, addition of cofilin, which promotes actin disassembly in conjunction with Cap32/34, abolished actin thickening in response to local administration of rapamycin. With Cap32/34 at a relatively low concentration (50 *nM*), we did not observe membrane protrusion or invagination even in the presence of higher concentrations of G-actin (Supplementary Figure 9), suggesting that shape deformations were bottlenecked by actin polymerization. Taken together, these results suggest that the timescale of ActA stability is critical to achieving a stable actin polymerization output, and that actin polymerization is needed for shape deformations. Additionally, these experiments further suggest that symmetry breaking is associated with local fluctuations in rapamycin concentration.

## Discussion

In vitro cell mimetic systems aim to assemble the minimal set of protein and lipid modules to output a desired function. Here, we have developed a cell mimetic platform that implements spatiotemporal control modules to achieve biological symmetry breaking with actin. To our knowledge, our platform comprises the most reduced network of biological nodes that is able to produce symmetry breaking in a context relevant to diverse physiological processes, including chemotaxis. Specifically, our design is devoid of capping protein, myosin, or phase-separated lipids that were previously considered indispensable to symmetry breaking.^24, 49, 50^ Using our synthetic platform, we have found that the application of undirected external chemical inputs results in directed actin polymerization and asymmetric membrane deformations. The actin polymerization and shape deformations are uncorrelated with upstream biochemical cues, indicating biological symmetry breaking. Biophysical modeling suggests that a model of actin polymerization is consistent with our experimental observations and can lead to substantive membrane shape changes through mechanical deformations. Local experiments, in which rapamycin is directionally administered, further confirm these dynamics and suggest that actin symmetry breaking arises from local fluctuations in rapamycin concentration.

Our work therefore shows that signal amplification from upstream chemical components, including Arp2/3, to actin is sufficient to drive symmetry breaking. Moreover, our platform exemplifies how implementing CID in an active model system can lead to changes in the system’s mechanical and morphological properties. We observed that seconds-long local ActA enrichment on the membrane close to the input was sufficient to bias the F-actin towards the input source, a finding which points to the potential positive feedback characteristic of actin polymerization in response to ActA-induced Arp2/3 activation^51^ and which could be further interrogated with Arp2/3 mutant variants for more fine-tuned control of symmetry breaking.

Our biophysical modeling of the dynamics and mechanics of actin polymerization underscores the interplay between actin symmetry-breaking and GUV shape deformations. Other models have been developed to describe actin dynamics and mechanics in other contexts; notably, these models have focused on different aspects of actin polymerization, including actin flow^22^ and settings of reduced dimensionality.^44^ The model developed in this work is specific to our platform in that it focuses on the patterns of actin polymerization generated through CID, as well as the stretching deformations caused by an internal osmotic pressure. Our model suggests that the mechanical stress-dependent incorporation of actin subunits at actin nucleation sites is consistent with our empirical observations, and that the coupling of asymmetric actin patterning with thin-shell mechanical properties alone can generate asymmetric GUV shapes. Our model and its conclusions may generalize beyond our experimental platform to offer physical insights into other processes, including actin-dependent cell shape changes and cell motility.

In general, bottom-up approaches that rely on de novo assembly to understand biophysical principles have recently gained traction in synthetic biology. Our work exemplifies these efforts on the molecular scale. The presented platform provides insight on the spatiotemporal localization of the molecular components and elucidates the design principles governing chemical signaling, actin assembly, and symmetry breaking. Our platform offers a novel GUV manipulation technique^52, 53^ and a versatile imaging pipeline to track the locality-specific concentration of biomolecules in a dynamic GUV over time, complementing other realized modalities.^54, 55^ Our developed image processing and statistical analysis scheme enables probing the biomolecular and geometric features of symmetry breaking, a pipeline that is is applicable to other GUV- and cell-based studies. By combining experiment and modeling, our results offer the ability to better understand macroscopic cellular processes, including morphogenesis, motility, and division, that build on fundamental biophysical and active matter principles to accomplish complex biological tasks.

## Methods

### DNA plasmid construction and protein purification

Detailed in Supplementary Materials.

#### Global CID-based protein translocation in GUVs

For CFP-FRB translocation towards the mCherry-FKBP-MARCKS-bound membrane, we reconstituted 4.4 μM FRB-containing and 7.8 μM FKBP-containing constructs in PBS buffer inside symmetric GUVs (Supplementary Table 1) using the inverted emulsion-based GUV fabrication technique previously reported (41). To balance the osmotic pressure, 750 *mM* sucrose was reconstituted inside while the GUVs were collected in 750 *mM* glucose containing PBS buffer. 100 μM DMSO-solubilized rapamycin was administered in the outer GUV buffer to trigger translocation. For actin polymerization experiments we reconstituted the components reported in Table 1.

#### Global rapamycin administration image acquisition

For CFP-FRB translocation experiments, we used an inverted epi-fluorescence microscope (Axiovert135TV, ZEISS) with 40× oil objective to track the CFP and mCherry signal intensity at one frame per 60 seconds. For actin polymerization experiments we used LSM780 confocal microscope (Zeiss) equipped with Plan-Aprochromat 63X/1.40 N.A. oil immersion DIC objective lens (Zeiss 420782-9900).

#### Image processing and biophysical parameters extraction

For Figure 1 and Supplementary Figure 1 MetaMorph® software (Molecular Devices) was used to measure the average luminal and membrane fluorescence intensity for either CFP or mCherry. Values are reported after normalizing by the average intensities of each prior to rapamycin or DMSO treatment. All actin-containing microscopy data was analyzed using a custom MATLAB® (MathWorks) script (Supplementary Material and code). For pre-processing, image sequences were regionally cropped and smoothed with 2D median and Gaussian filtering. Individual GUVs were segmented based on membrane marker signal (mCherry-FKBP-MARCKs) using intensity thresholding, morphological operations, and active contours technique.

#### Local rapamycin administration with a microinjector and image acquisition

The GUV content is listed in Supplementary Table 2. A 100 μL droplet of freshly prepared GUVs was placed in a 1-well Lab-Tek II chambered cover glass (Thermo Scientific) immediately before imaging. Local stimulation of GUVs was carried out by lowering a micropipette (Femtotips, Eppendorf) loaded with 10 μL of a freshly prepared solution of 500 μM rapamycin and 100 μM AlexaFluor 647 dye (Thermo Scientific) and applying microinjection near GUVs of interest. The rapamycin-dye solution was rapidly prepared on ice in a 100 μL volume, diluted in double distilled H_2_O from stock solutions of rapamycin (5*mM* in 100% ethanol) and AlexaFluor 647 (1 *mM* in DMSO). Concentrations were optimized based on estimates of micropipette-generated chemical gradients under passive diffusion or forced flow^56^ and considerations of needle clogging. The micropipette was then connected to a microinjector (Femtojet, Eppendorf) controlled by a micromanipulator (Eppendorf). Image acquisition began with the micropipette immersed in the liquid droplet but above the focal plane of GUVs, at an initial compensation pressure (*P*_*c*_) of 5-10 hPa. To achieve local chemical dimerization, the micropipette was lowered near a GUV in the field of view, and compensation pressure was increased to between 30 – 50 hPa for a continuous local gradient. To relieve clogging and generate brief local gradients, microinjection pressures were also applied in bursts as needed (injection parameters: injection pressure (*P*_*i*_) = 400 - 800 hPa; injection duration (*t*_*i*_) = 5-10 sec; *P*_*c*_ = 10 hPa). Confocal image acquisition every 15 seconds was performed on LSM780 confocal laser-scanning microscope (Zeiss).

#### Reagents

Rabbit skeletal muscle actin (AKL99), pyrene-labeled actin (AP05), Arp2/3 complex (RP01P), and GST-WASP-VCA (VCG03-A) were purchased from Cytoskeleton. ATP for actin dialysis buffer was purchased from Gold Biotechnology. All lipids (Supplementary Table 1) were purchased from Avanti Polar Lipids. Hexadecane (H6703) and silicone oil (viscosity 20 cSt (25 °C), 378348) were purchased from Sigma Aldrich.

#### Statistical Analysis

Symmetry breaking analysis from kymographs using correlation analysis, and principal component analysis are detailed in Supplementary Materials. To verify our method, we applied our analysis pipeline to a previously reported case of symmetry breaking in GUVs (Supplementary Figure 12).^43^ Please refer to the figure legends for the description of sample size and the corresponding statistical methods used.

## Acknowledgements

We thank Dr. Tianzhi Luo for assistance with vesicle fabrication techniques and Elmer Rho for help with cloning and analyzing FRET data. We thank Dr. Cécile Sykes, Dr. Winston Timp, and Dr. Rong Li for invaluable discussions, Dr. Erin Goley for suggestions and support of the FPLC and fluorimetry experiments, Dr. Daniel Raben for sharing lipid handling setup, and Dr. Koki Kamiya and Dr. Masashi Tabuchi for experimental suggestions on lipid membranes and GUV fabrication. We thank Dr. Masato Kanemaki for the strep-tag plasmids and Robert DeRose for proofreading the manuscript. We are also grateful to Dr. Cristina Martinez-Torres group for generously sharing the beta version of the DisGUVery image analysis pipeline with us.

## Data and materials availability

All the data associated with this work are presented in the manuscript and its associated supplementary files.

## Code availability

General MATLAB® code for use in .tif file processing and analysis are available under an open-source license in the GitHub repository https://gitfront.io/r/basharif/gTdvogxUoH4y/GUV-symmetry-breaking-analysis/. Specific .m scripts for each experiment, as well as Abaqus FEA input files used for finite-element simulations, are available from the corresponding authors upon request.

## Author contributions

S.R. and T.I. conceived the project. S.R. designed and performed the experiments and analyzed data. S.R. implemented the microinjector experiments with help from B.A.-S. F.W. performed modeling. B.A.-S. developed the image processing and statistical analysis pipeline with input from P.I., F.W., and S.R. H.T.M. performed the pyrene assay and purified and labeled actin with help from B.C. H.N. developed and characterized the ActA tool. P.I., J.L., and E.S. helped with modeling. D.N.R. provided protein purification and GUV fabrication resources. S.R. and F.W. wrote the manuscript with help from T.I., and B.A.-S. All authors contributed to editing of the manuscript.

## Funding

This work was supported by the National Institute of Health (5R01GM123130, R01GM136858, R35GM149329 to T.I., R35GM128786 to B.C., and S10OD016374 to Scot C. Kuo), the DoD DARPA (HR0011-16-C-0139 to P.A.I., D.N.R. and T.I.), the National Science Foundation (000819255 to T.I., J.L., B.C.), and the PRESTO program of the Japan Science and Technology Agency (PRESTO, JPMJPR12A5 to T.I.). S.R. was supported by the National Science Foundation and the Japan Society for the Promotion of Science (JSPS) East Asia and Pacific Summer Institutes and Johns Hopkins Lucille Elizabeth Hay graduate fellowships.

## Competing interests

S.R., T.I., D.N.R. are inventors on a filed GUV fabrication patent (US2021/0145746 A1).

## Notes

### Competing Interest Statement

The authors have declared no competing interest.

