## Supplemental Material for "Synthetic control of actin polymerization and symmetry breaking in active protocells"

**This PDF file includes:**

- Supplementary Text
- Supplementary Figures 1 to 12
- Supplementary Tables 1 to 2
- Supplementary Movies 1 to 3 Captions

**Other Supplementary Materials for this manuscript include the following:**

- Supplementary Movies 1 to 3

### Supplementary Text

#### Plasmid construction

Twin-Strep-tag®-pET28 was a kind gift from Dr. Kanemaki. pQE-80L-based plasmids and SspB sequences were obtained from pQE-80L MBP-SspB Nano (Addgene #60409). Mammalian expression plasmids and ECFP, EYFP, and mCherry sequences were constructed and obtained from pECFP-C1, pEYFP-C1, and pmCherry-C1 (Clontech). The mActA(1-183) sequence was PCR- amplified from the plasmid reported previously.[59] The beta-actin sequence was obtained from paGFP-actin plasmid, as previously reported (80). MARCKS and Fyn sequences were synthesized by oligo DNA annealing.

#### Protein expression and purification

**6xHis-YFP-FKBP (pET28a)** We transformed YFP-FKBP in BL21 (RIL) cells and set up an overnight, 37 $^{\circ}$C, 220 rpm culture of one colony in 5 mL of LB media supplemented with kanamycin and chloramphenicol. The next day, we transferred the 5 mL culture to 1 L of LB media supplemented with kanamycin and chloramphenicol at the same growth condition. Cells were induced at $OD_{600}$ of 0.5 with 400 $\mu M$ IPTG, and the temperature was reduced to 19 $^{\circ}$C for overnight growth. Cells were harvested the next day, resolubilized in lysis buffer (10 $mM$ Tris-HCl pH 7.4, 50 mM NaCl, 10 $mM$ imidazole, 10 mM β-mercaptoethanol, 10% V/V glycerol), and lysed with the microfluidizer. Lysate was spun at 20,000x g for 40 min to isolate the soluble fraction, and cleared with 0.2 $\mu m$ filter. The soluble fraction was loaded on Ni-NTA beads (GE Healthcare) and washed with 15 column volumes wash buffer (10 $mM$ Tris-HCl pH 7.4, 500 mM NaCl, 10 $mM$ imidazole, 10 mM β-mercaptoethanol, 10% V/V glycerol). An FPLC was used to administer an imidazole linear gradient at 200 $mM$ imidazole peak (10 mM Tris-HCl, 100 $mM$ NaCl, 200 $mM$ imidazole, 10 $mM$ β-mercaptoethanol, 10% V/V glycerol). The purest fractions (assessed by SDS-PAGE gel) were harvested and dialyzed overnight in storage buffer (10 $mM$ Tris-HCl pH 7.6, 50 $mM$ NaCl, 10% V/V glycerol, 1 $mM$ DTT). Protein was concentrated (Amicon Ultracentrifugation), snap frozen, and stored at -80 $^{\circ}$C.

**6xHis-CFP-FRB (pET28a)** We transformed CFP-FRB in BL21 (RIL) cells and inoculated one colony in 5 mL of LB supplemented with kanamycin media and grew at 37 $^{\circ}$C, 220 rpm, overnight. 5 ml of the overnight culture was added to 1L of LB/kanamycin media and induced with 400 $\mu M$ IPTG when $OD_{600}$ reached 0.5. Growth was continued at 19 $^{\circ}$C overnight and cells harvested the next day, resolubilized in lysis buffer, and lysed with a microfluidizer. Lysate was spun at 20,000$g$ for 40 min and the soluble fraction collected. To pellet down DNA, 5% W/V polyethyleneimine was gradually added and spun at 15,000$g$ for 5 minutes. The supernatant was collected and saturated ammonium sulfate added for a final concentration of 40%, spun at 10,000$g$ for 10 minutes. Pellet was resuspended in storage buffer. We then dialyzed the protein overnight to remove excess ammonium sulfate and used Ni-NTA column for the remaining purification steps similar to what is detailed for YFP-FKBP construct.

**6xHis-mCherry-FKBP (pBiEx) and 6xHis-mCherry-FKBP-MARCKS (pBiEx)** BL21-RIL cells were transformed. 10 ml of overnight LB/ampicillin culture was grown at 37 $^{\circ}$C and 220 rpm was transferred to 1L of the same media for next day expression at 37 $^{\circ}$C. Due to high level of leaky expression, no further IPTG induction was implemented. The culture was harvested after 15-20 hours of growth when the media color looked brightly pink. Cells were resolubilized in the lysis buffer (50 $mM$ Tris-HCl pH 7.9, 100 $mM$ NaCl, 1 mM EDTA, 10% glycerol, 10 $mM$ β-Mercaptoethanol, Roche EDTA-free protease inhibitor), lysed with the microfluidizer, spun at 20,000$g$ for 40 min and the soluble fractions were harvested. Ni-NTA purification proceeded similar to what is detailed for YFP-FKBP.

**6xHis-mActA (1-584)-FRB-CFP (pET28a)** We inoculated a colony of BL21-CodonPlus(DE3)-RIL competent cells (Agilent Technologies 230245) transformed with the plasmid DNA in 10 ml of LB media supplemented with kanamycin and grew overnight at 37 $^{\circ}$C, 220 rpm. Culture was transferred to 1L of LB/kanamycin and expanded at 37 $^{\circ}$C, 220 rpm till $OD_{600}$ reached 0.5. Cells were induced with IPTG at a final concentration of 0.4 $mM$ and culture continued at 19 $^{\circ}$C, 220 rpm, overnight. The cells were pelleted at 4500$g$, 4 $^{\circ}$C for 15 min and the pellet resolubilized in 20 ml of lysis buffer (10 $mM$ Tris pH 7.6, 100 $mM$ NaCl, 10 $mM$ imidazole, 10% glycerol, 1 $mM$ β-mercaptoethanol and cOmplete™ EDTA-free protease inhibitor tablet (Sigma-Aldrich 11836170001). Cells were lysed with a microfluidizer and the soluble fraction harvested by centrifugation at 16,000$g$, 4 $^{\circ}$C for 40 min. The supernatant was filtered with 0.2 $\mu m$ syringe filters (Thermo Scientific™ F25006). We equilibrated a 5 ml His-Trap HP column (GE Life Sciences) with the lysis buffer lacking protease inhibitors and eluted the protein using a linear imidazole gradient from 50-500 $mM$ over 20 column volumes. The eluted fractions were analyzed on a SDS-PAGE gel and subsequently the peak fractions detected on the Coomassie blue (Biorad 1610400) stained gel were pooled and dialyzed in a 10 kDa SnakeSkin™ dialysis tubing (Thermo Fisher Scientific 68100) in 2L of storage buffer (10 $mM$ Tris-HCl pH 7.6, and 50 $mM$ NaCl, 10% glycerol, and 1 $mM$ β-mercaptoethanol) at 4 °C, overnight. The next day the protein was dialyzed for an additional 3 hours in fresh storage buffer and subsequently applied to a Superdex 200 10/300 GL (GE Life Sciences) equilibrated in the same buffer. The peak fractions were combined, and concentrated with an Amicon® Ultra 10 kDa centrifugal filter unit (Millipore UFC801024). Aliquoted protein vials were flash-frozen and stored at -80 °C.

**2xstrep-ActA(1-183)-FRB-CFP (Twin-Strep-tag®-pET28)** Inoculation and growth was similar to the ActA(1-584)-CFP-FRB construct. For lysis we used 100 $mM$ Tris-HCl pH7.4, 150 $mM$ NaCl, and cOmplete™ EDTA-free protease inhibitor. The cells were lysed with a microfluidizer, and the supernatant clarified as previously detailed. Strep-tactin XT (GE Healthcare) beads were equilibrated with 100 $mM$ Tris-HCl, 150 $mM$ NaCl, 1 $mM$ EDTA, pH 8.0 and the cleared lysate was passed through the column and washed with 10 column volumes of the same buffer. Protein was eluted with 100 $mM$ Tris-HCl, 150 $mM$ NaCl, 1 $mM$ EDTA, and 50 $mM$ biotin at differential elution volumes recommended by the manufacturer, dialyzed against storage buffer (10 $mM$ Tris-HCl, 50 $mM$ NaCl, 10% glycerol, 1 $mM$ DTT) overnight, concentrated, and froze down in individual vials the next day.

**Actin** One milligram of actin (Cytoskeleton Inc. AKL99-B) was dissolved in 400 µL G-buffer (2 $mM$ Tris-HCl (pH 7.5 at RT), 0.1 $mM$ CaCl2, 0.2 $mM$ ATP, 0.5 $mM$ DTT, 1 $mM$ $NaN_3$) and dialyzed against the same buffer. After 3 days dialysis with daily buffer exchange, the actin was purified by Superdex 200 Increase 10/300 GL column (GE Healthcare) to separate monomer fraction from occasional larger size (smaller elution volume) fraction. The purified actin was stored at 4 °C and dialyzed against the G-buffer that was refreshed twice a week.

**Cap32/34 (pET14b)** Cap32 and Cap34 proteins were individually expressed in LB/Amp culture (conditions similar to CFP-FRB). The pellets were pulled together after resolubilization in the lysis buffer (used for CFP-FRB) to form the Cap32/34 complex. Ni-NTA column was used for purification followed by size exclusion column to isolate the protein complex fractions.

#### FRET assay in the GUVs

3 $\mu M$ CFP-FRB and 3.4 $\mu M$ YFP-FKBP were reconstituted in PBS buffer inside symmetric GUVs with POPC membrane. For osmotic pressure balance fsupp 750 $mM$ sucrose was present in the GUVs while outside buffer was PBS buffer with 750 $mM$ glucose. An inverted epi-fluorescence microscope (Axiovert135TV, ZEISS) with 40× oil objective was used to take images at one frame per 30 seconds prior to and post-rapamycin and DMSO (control) administration. CFP, YFP, and FRET intensities were measured. The FRET signal was reported after normalizing against the CFP donor signal. Image processing was performed using the MetaMorph® software (Molecular Devices). To account for photobleaching, the decay in fluorescence signal 10 min prior to input addition was measured and the data were corrected approximating photo bleaching as a linear trend.

#### Pyrene assay

Pyrene-actin polymerization assay was performed with FluoroMax 3 and Datamax software. Pyrene fluorescence and its kinetics were measured by 365 $nm$ emission (1 $nm$ bandwidth) and 497 $nm$ excitation (5 $nm$ bandwidth). Actin and pyrene-labeled actin were mixed in G-buffer to make 5% pyrene actin. Arp2/3 and GST-WASP-VCA were reconstituted as described in the manufacturer's protocols and diluted to desired concentration with F-buffer (10 $mM$ Tris-HCl (pH 7.5 rt), 50 $mM$ KCl, 2 $mM$ MgCl2, 1 $mM$ ATP). Reaction mixture was prepared as indicated in figure legends and the reaction was performed at room temperature. Reaction was started with 1 $\mu M$ actin and 10 $nM$ Arp2/3 solution in F-buffer. GST-WASP-VCA or ActA variants (final 100 $nM$) were added at time = 0. Reaction was started with 1 $\mu M$ actin in F-buffer. Arp2/3 (final 10 $nM$) and GST-WASP-VCA or ActA variants (final 800 $nM$) were added at time = 0.

#### Image processing and analysis

**Kymograph generation** To generate the kymographs, an initial list of finely and equally spaced boundary points were computed using a snake algorithm for each frame based on the segmentation mask. A 2-D Savitzky-Golay filter was applied to generate a smooth boundary for estimating biophysical boundary quantities. Biochemical quantities (membrane marker, ActA, actin) were estimated at each boundary point as the mean of the background-subtracted fluorescence intensity along a short line segment perpendicular to the boundary. For the kymographs used in PCA analysis, biochemical quantities were quantified as boundary:lumen ratios by normalizing the fluorescent signal at a given boundary point (computed as before) to the background-subtracted fluorescent signal in the lumen (computed as background-subtracted mean of the bottom 20% of signal values in a mask defining the GUV lumen). Local curvature was approximated as 1/radius of the circle with best geometric fit for a local set of boundary points (code from MATLAB file exchange). To generate final kymographs with a fixed number of points for all subsequent analysis, the list of biophysical quantities initially estimated over the smooth, dense boundary points was sub-sampled at 360 equally spaced angular bins from $-\pi$ to $\pi$.

**Correlation analysis** For tracking correlations between local biophysical quantities in individual GUVs over time, the Pearson correlation coefficient was computed for pairs of biophysical quantities (ActA-actin; ActA-curvature; actin-curvature) over 360 boundary points at each frame (a single column in a kymograph). To minimize the effect of artificial correlations due to membrane artifacts, an inclusion criteria for both spatial points and frames was used: 1) only include spatial points with membrane marker signal in bottom 75th percentile for a given GUV, and 2) only include frames with at least 180 points. All available GUVs (n=18) were utilized in this analysis.

The evolution of asymmetry in biochemical parameters and global shape changes in individual GUVs over time were assessed using two quantities: the entropy of biochemical signal values, and the eccentricity of the GUV. Changes in entropy, computed in the information theory sense, as a measure of signal randomness, provided an indirect measure of changes in symmetry. The input space of biochemical signals from all frames of an individual GUVs kymograph was first discretized into 256 bins for consistent 8-bit representation. For each GUV, the entropy for a given biochemical quantity at the jth frame, $X^{(j)}$ , was then computed in the information theory sense as $H(X^{(j)}) = \sum_i p_i log_2(p_i)$, for all $p_i > 0$, where $p_i = P(X^{(j)} = x_i)$ is the probability of the ith bin of signal values. The probability distribution for $X^{(j)}$ was computed from the histogram of discretized signal values at frame $j$. Fixed spatial discretization at each frame (360 spatial points) and fixed signal discretization (8-bit) ensured consistency with entropies remaining between 0 and 8 over all frames and all GUVs. For tracking global shape at each frame, MATLAB's built-in function was used to compute eccentricity, defined as the ratio of the focal length to the major axis length of the ellipse that best fits the GUV mask. Note that eccentricity values range from 0 (perfect circle) to 1 (line segment). All available GUVs (n=18) were utilized in this analysis.

To compare fluorescent intensity profiles of ActA and actin from different GUVs before and after rapamycin, two time points were selected: two frames before rapamycin addition, and 10 frames after rapamycin addition. At each frame, the mean-normalized fluorescent intensity profiles of ActA and actin across 360 spatial points were plotted. All available GUVs (n=18) were utilized in this analysis.

Bulk correlation analysis was used to assess biophysical trends in data from multiple GUVs (n=17) and multiple frames. Initially, biochemical kymographs were max-normalized for proper comparisons across GUVs. For ActA-actin bulk correlation analysis, the kymographs were split into a pre-rapamycin section (frame 1 to last frame before rapamycin; length varies by GUV but ranges from 6-18 frames) and a 35-frame post-rapamycin section (10 frames after rapamycin to 44 frames after rapamycin). Pearson correlation coefficients were computed for pairs of ActA and actin values in these sections from multiple GUVs. To minimize the effect of artificial correlations due to membrane artifacts, an inclusion criteria for spatial points was used: only include points with membrane marker signal in bottom 75th percentile for a given GUV.

For ActA-eccentricity and actin-eccentricity bulk correlation analyses, the mean of the biochemical signals from 360 spatial points was first computed at each frame. This scalar conversion of biochemical signals was necessary for comparison with the eccentricity, a scalar shape parameter. Additionally, all 360 points were included for scalar conversion; no inclusion criteria based on membrane marker was employed. Pre-rapamycin sections for each GUV were defined as before; however, post-rapamycin section was defined as a 35-frame window from 20 frames after rapamycin to 54 frames after rapamycin. Pearson correlation coefficients were then computed for pairs of biophysical quantities (mean ActA-eccentricity; mean actin-eccentricity) in these sections and multiple GUVs.

**Principal component analysis** Prior to applying principal component analysis (PCA) analysis on kymographs from multiple GUVs, steps were taken for signal normalization and spatiotemporal alignment. For ActA and actin, boundary:lumen ratio based kymographs were used, with max-normalization for each GUV. For curvature kymographs, the local curvature was normalized to an approximate bulk initial curvature for each GUV. This bulk initial curvature was approximated as $\frac{1}{0.5L}$ where L is the major axis length of the best-fit ellipse for the GUV shape at the first frame after rapamycin. Spatial points from all kymographs were binned from 360 points to 180 points using averaging. Smoothing was performed using a 21-point moving average window in space. Spatial alignment of kymograph data from globally stimulated GUVs (n=17) was approached by first centering each GUV's actin kymograph to the spatial point (corresponding to an angular direction) with maximum value at 35 frames post-rapamycin. This centering direction was then applied to the ActA and curvature kymographs for comparison of patterns across biophysical parameters. Temporal alignment was performed by simply selecting a 50-frame window from frame 1 post-rapamycin to frame 50 post-rapamycin. For locally stimulated GUVs (n=3), all kymographs were spatially aligned and centered to the direction of the external micropipette; and temporal alignment was performed by selecting a 250-frame window, from frame 1 post-rapamycin to frame 250 post-rapamycin.

PCA was then performed using singular value decomposition of the matrix of mean-subtracted, aligned, vectorized 2-D kymographs. This matrix has 17 columns for the global rapamycin data and 3 columns for local rapamycin data, from the 17 GUVs and 3 GUVs analyzed respectively for each dataset. Principal components (PCs) were re-shaped into the dimensions of the 2-D kymographs for visualization of dominant spatiotemporal trends in kymographs. Cumulative sums of the square of the singular values were used to compute percent variance captured by the corresponding top PCs.

#### Model of actin polymerization dynamics

As discussed in the main text, we assume that, due to baseline levels of spontaneous (rapamycin-independent) actin polymerization, there already exists a spherical shell layer of actin anchored to the inner leaflet of the vesicle.[60] This actin shell will be the main load-bearing element, and will be remodeled by additional (rapamycin-dependent) polymerization (Fig. 4a of the main text). While characteristics including actin flow[22] may also be important, here we focus on the elastic stretching stresses in the actin shell that are generated by the outward turgor pressure.

Suppose there is a nucleation site (a subsurface of the sphere) at which increased actin polymerization occurs. Any point on this site is given by surface coordinates, $(\theta, \varphi)$, where $\theta \in [0, 2\pi]$ and $\varphi \in [0, \pi]$ span the sphere. We assume that the volume, $V$, of polymerized actin is increased locally at this nucleation site. As described in the main text, we may consider a model in which the local, volumetric rate of actin polymerization at any coordinate $(\theta, \varphi)$ contained within the nucleation site quantitatively depends on the mechanical stress:

$$\frac{dV(\theta, \varphi, t)}{dt} = \ell^3 v_0 e^{\sigma(\theta,\varphi,t)/\sigma_0}. \tag{2}$$

Here, $\ell^3$ is the volume of an effective unit actin insertion with edge length $\ell$, $v_0$ is a baseline actin polymerization rate with units of 1/min (which may depend on upstream nucleation factors such as Arp2/3 and Barb), and $\sigma_0$ is a baseline value of mechanical stress at the nucleation site. Eq. (2) is the basic assumption of our model; namely, that actin polymerization at the nucleation site is spatially coupled to mechanical stress, and more actin polymerization occurs at regions of larger tensile stress. We note, in turn, that mechanical stress is trivially coupled to actin polymerization because the shell thickness changes as $dh(\theta, \varphi, t)/dt = \ell^{-2} dV(\theta, \varphi, t)/dt$: since we expect the mechanical stress to scale as $pr/h$, where $r$ is the radius and $h$ is the thickness, there is less mechanical stress at regions of larger thickness. We note that Eq. (2) makes an assumption similar to Eq. (3) in ref.,[44] in which the authors assumed that the actin polymerization rate outside a droplet, with actin polymerizing in only one direction, depends on the normal stress in the droplet: the functional form of the stress-dependence is a Kramer's rate. The main difference with respect to our Eq. (2) is that Eq. (2) considers the in-plane, not normal, stresses, and that actin polymerization will occur in three dimensions.

In Fig. 4 of the main text, we have plotted $\sigma$ as the maximal principal stress of the actin shell as a simple scalar measure of tension:

$$\sigma_{\mathsf{M}} = \max(\sigma_{xx}, \sigma_{yy}). \tag{3}$$

where $\sigma_{xx}$ and $\sigma_{yy}$ are the nonvanishing components of the stress tensor in the orthogonal coordinate system $(x, y)$. Nevertheless, it is straightforward to see that our results hold for other stress measures.

We will show that, with the assumption of Eq. (2), the model predicts the lateral and normal modes of actin polymerization, and also the observed vesicle shrinkage. We first estimate the mechanical stress, $\sigma(\theta, \varphi, t)/\sigma_0$.

##### Determination of the mechanical stresses in an axisymmetric polymerizing actin shell

We first determine the mechanical stress in the polymerizing actin shell. This analysis will reveal that, when actin polymerizes locally at a nucleation site, the mechanical stresses at the nucleation site are largest at the edges. Accordingly, from Eq. (2), the direction of fastest polymerization will be lateral on the surface of the vesicle. Furthermore, we will quantitatively determine the magnitude of stress increase, which will dictate the value of the $e^{\sigma(\theta,\varphi,t)/\sigma_0}$ term in Eq. (2) and allow for the estimation of $v_0$ from experimental data.

We assume that polymerized actin forms a linear-elastic, isotropic, spherical shell anchored to the inner leaflet of the vesicle. Denote the radius, thickness, three-dimensional Young's modulus, and Poisson's ratio of this shell as $r$, $h$, $E$, and $\nu$, respectively. This shell resists an internal (osmotic) pressure of $p$. If the shell thickness, $h$, is constant, then we know from thin shell theory that the stresses, $\sigma$, and strains, $\varepsilon$, in an orthogonal coordinate system, $(x, y)$, are

$$\sigma_{xx} = \sigma_{yy} = \frac{pr}{2h}, \quad \sigma_{xy} = 0, \quad \varepsilon_{xx} = \varepsilon_{yy} = \frac{\sigma}{E}(1-\nu) = \frac{pr(1-\nu)}{2Eh}, \quad \varepsilon_{xy} = 0. \tag{4}$$

The introduction of a nucleation site results in actin polymerization on a patch of this shell. This increases the actin thickness, $h$, locally, so that $h$ is no longer a constant but varies in space and time. Let us assume, for simplicity, that actin polymerization increases $h$ axisymmetrically (independent of $\theta$) beyond a baseline value, $h_0$, by a constant fold amount, $f$. Under the assumption of constant $f$, $h(\theta,\varphi,t) = h(\varphi,t)$ depends only on the subtended angle $\varphi$ as follows:

$$h(\varphi,t) = \begin{cases} fh_0 & \varphi < \varphi_0(t) \\ h_0 & \varphi \geq \varphi_0(t). \end{cases} \tag{5}$$

In particular, the spherical shell transitions from a thickness of $fh_0$ to a thickness of $h_0$ at the angle $\varphi_0(t)$, and this angle increases in time; ultimately, if actin polymerizes faster laterally than in the thickness direction, the entire shell will have thickness $\approx fh_0$ due to the lateral actin polymerization.

In the bulk of the two caps ($\varphi \ll \varphi_0(t), \varphi \gg \varphi_0(t)$), we expect analogous equations to Eq. (4) to hold because the thickness change is only at the interface. The mechanical stress at the interface may be more subtle because of this thickness change. We anticipate that the stresses will increase from $\sigma_{xx} = \sigma_{yy} = pr/2fh_0$ to $\sigma_{xx} = \sigma_{yy} = pr/2h$ at the interface. In confirmation of this, we used finite-element simulations (Abaqus FEA) to model the stresses at the interface. Simulating the mechanical stresses across different values of $f$ and $\varphi_0$, the results consistently revealed that the stresses at $\varphi = \varphi_0$ increases from the stresses in the thicker region to the stresses in the thinner region (Fig. 4b of the main text). Based on these results, and normalizing with respect to the baseline stress $\sigma_0 = pR/(2fh_0)$ at the nucleation site, we find that for $\varphi \leq \varphi_0$,

$$\frac{\sigma(\theta,\varphi,t)}{\sigma_0} \approx \begin{cases} 1 & \varphi < \varphi_0(t) \\ f & \varphi = \varphi_0(t). \end{cases} \tag{6}$$

In other words, the fold-change in stress is maximal at the edge of the nucleation site and is approximated by the fold-change, $f$, in actin thickness. $f$ can be measured from the experiments by comparing the thickness from the actin fluorescence before and after lateral actin polymerization.

#### Determination of the lateral and normal actin polymerization rates

Based on the considerations above, the exponential term in Eq. (2) will be constant in time at the expanding front ($\varphi = \varphi_0$), and the term will be approximately $e^f$. Thus, for $f \gg 1$, the model predicts that actin polymerization will occur mostly laterally whenever there is differential stress in the actin shell. When the actin shell becomes uniform in thickness (under our simplifying assumptions, with a thickness of $fh_0$), there is no differential stress and $\sigma(\theta,\varphi,t)/\sigma_0 = 1$; accordingly, the polymerization is expected to occur in the normal, out-of-plane direction, and its rate decreases by a factor of $e^{f-1}$. This can be summarized as follows:

$$\frac{dV(\theta,\varphi,t)}{dt} \approx \begin{cases} \ell^3 v_0 e^f & \varphi = \varphi_0(t) \\ \ell^3 v_0 e & \varphi < \varphi_0(t). \end{cases} \tag{7}$$

Using this equation, we can determine how the actin circumference and thickness should increase in time.

During lateral actin polymerization, the area that is added laterally (at $\varphi = \varphi_0$) to the actin shell is added at any angle $\theta$ with rate

$$\frac{dA(\theta,t)}{dt} = \frac{1}{\ell}\left.\frac{dV(\theta,\varphi,t)}{dt}\right|_{\varphi=\varphi_0} \approx \ell^2 v_0 e^f, \tag{8}$$

so the total lateral polymerized area at $\theta$ after time $t$ is

$$A(\theta,t) = \ell^2 v_0 e^f t. \tag{9}$$

The total area of an axisymmetric spherical cap of depth $d$ is $2\pi rd$. Integrating $A(\theta,t)$ over $\theta \in [0,2\pi]$ and setting the two areas equal, the total polymerized depth after time $t$ is $d(t) = \ell^2 v_0 e^f t/r$. The angle subtended by the axisymmetric spherical cap is $\varphi_0(t) = \cos^{-1}[(r-d)/r]$. Taking $r$ to be constant and approximated by $r_0$ (which is valid for small deformations and for the short timescale of lateral actin polymerization), the total polymerized circumference at time $t \le \varphi_0^{-1}(\pi)$ is then

$$C(t) = C_i + 2\pi r \times \frac{\varphi_0(t)}{\pi} = C_i + 2r\cos^{-1}\left(1 - \frac{d(t)}{r}\right) = C_i + 2r\cos^{-1}\left(1 - \frac{\ell^2 v_0 e^f t}{r^2}\right), \tag{10}$$

where $C_i$ is a constant initial value (and will hereafter be taken as 0). Note that $\varphi_0^{-1}(\pi)$ is the time at which lateral polymerization completes, and the entire circumference is covered with the laterally polymerized actin.

Although most of the actin will polymerize laterally (under the assumption that $f \gg 1$), there is also normal, out-of-plane polymerization in the bulk of the nucleation site ($\varphi < \varphi_0$). The volume that gets added to the actin shell at any coordinate $(\theta,\varphi)$, $\varphi < \varphi_0(t)$, is added with rate

$$\frac{dV(\theta,\varphi,t)}{dt} \approx \ell^3 v_0 e. \tag{11}$$

Integrating over $\theta \in [0,2\pi]$, $\varphi \in [0,\varphi_0(t)]$, and $t$, the volume added by this mode of polymerization after time $t$ is

$$2\pi\ell^3 v_0 e \int_{t'=0}^{t} \cos^{-1}\left(1 - \frac{\ell^2 v_0 e^f t'}{r^2}\right) dt',$$

which can be calculated numerically. Distributing this volume over the subtended surface area of $2\pi rd$, the average thickness of the laterally polymerized actin increases as

$$h(t) = h_i + \frac{\ell}{e^{f-1}t}\int_{t'=0}^{t} \cos^{-1}\left(1 - \frac{\ell^2 v_0 e^f t'}{r^2}\right) dt', \quad t \le \varphi_0^{-1}(\pi), \tag{12}$$

where $h_i = fh_0$ is a constant initial value. For $f \gg 1$, the second term becomes vanishingly small and is dominated by the first term. Thus, the thickness increase does not significantly increase the ratio $h(t)/h_0$ beyond $f$ during lateral polymerization, indicating the self-consistency with assuming $f$ to be constant in the limit of large $f$.

Similarly, during the normal, out-of-plane polymerization which occurs after the actin shell has uniform thickness, $\varphi_0 = \pi$, $d = 2r$, and the average thickness of the polymerized actin increases as

$$h(t) = h_i' + \frac{\pi\ell^3 v_0 e t}{2r^2}, \quad t > \varphi_0^{-1}(\pi), \tag{13}$$

where $h_i' = h(\varphi_0^{-1}(\pi))$ is a constant initial value.

#### Determination of the radial shrinkage rate during actin thickening

As actin polymerizes uniformly, the thickness of the actin shell increases as given by Eq. (13). The strain is given by Eq. (4) and decreases in time:

$$\varepsilon(t) = \frac{pr(1-\nu)}{2Eh(t)} = \frac{pr(1-\nu)}{2E(h_i' + \pi\ell^3 v_0 e t/(2r^2))} \approx \frac{pr_0(1-\nu)}{2E(h_i' + \pi\ell^3 v_0 e t/(2r_0^2))}. \tag{14}$$

In the above, we have assumed that $r$ is constant and approximately $r_0$, which is valid for small deformations and on the short timescale of lateral polymerization; deviations from this assumption will lead to minor corrections in the strain, which can be discarded for our parameter values of interest. Accordingly, with the typical linear strain-displacement relation, the model predicts the rate of shrinkage as:

$$\frac{r(t)}{r_0} = 1 + \varepsilon(t). \tag{15}$$

##### Comparison to experimental data

The main predictions of our model are the following:

$$\frac{C(t)}{2\pi r} = \frac{1}{\pi}\cos^{-1}\left(1 - \frac{\ell^2 v_0 e^f t}{r_0^2}\right), \quad h(t) = h_i' + \frac{\pi \ell^3 v_0 e t}{2r_0^2}\ (t > \varphi_0^{-1}(\pi)), \quad \frac{r(t)}{r_0} = \frac{p r_0 (1-\nu)}{2Eh(t)}. \tag{16}$$

- The first equation describes the time-evolution of the fraction of the vesicle contour that is covered by actin polymerizing on a nucleation site.
- The second equation describes the time-evolution of the average actin thickness. At times longer than the time it takes for the actin to extend around the circumference $(t > \varphi_0^{-1}(\pi))$, the thickness growth is linear in time, and actin polymerizes in the normal direction, uniformly over the entire surface.
- The third equation describes the radial shrinkage associated with the increase in thickness.

We anticipate that all parameters can be measured from the experiments, and that the only fitting parameter is $v_0$, the baseline rate of actin polymerization (with units of min$^{-1}$).

To compare with the experiments:

- We measured the thickness (in $\mu$m) across which actin fluoresces at the vesicle contour, and found that the fold-change in actin thickness during lateral polymerization was $f \gtrsim 2$. After a point on the vesicle circumference is covered by the laterally polymerizing actin, the thickness further increases slightly (order of $\sim$0.1-1$\times$) during this first phase of polymerization.
- We measured the vesicle radius, $r$, from either the phase-contrast or fluorescence (when phase-contrast not available) timelapse. We then used a linear approximation for the lateral polymerization rate as $2\pi rs/t_{\text{cir}}$, where $s \in [0, 1]$ is the fraction of the circumference polymerized and $t_{\text{cir}}$ is the time (min) needed to finish lateral polymerization (estimated in our model by $\varphi_0^{-1}(\pi)$).
- We measured actin thickness, $h$, across time in the fluorescence timelapses. We then used a linear approximation for the rate of thickness increase as $\Delta h/\Delta t$, where $\Delta h$ is the difference in actin thickness between the end of the timelapse ($t_{\text{final}}$) and the end of lateral polymerization ($t_{\text{cir}}$).
- Finally, we measured the radial shrinkage to be the fractional decrease in radius between the initial and final times.

For comparison, the model predictions for characteristic parameter values are shown in Fig. 4c of the main text. In particular, based on our empirical observations we assume that $f = 2$, $h_i = \ell = 0.1\ \mu$m, $r_0 = 20\ \mu$m.

- We estimate $v_0$ by approximating the lateral expansion velocity shown in the empirical data in Fig. 4c of the main text, which suggests $v_0 \approx 200$ min$^{-1}$: this value predicts that actin polymerization would cover the vesicle contour after $\varphi_0^{-1}(\pi) \sim$ 15 min, a timescale consistent with experiments.
- Over a longer timescale of $\sim$100 min, $h(t)$ is predicted to increase $\sim$10-fold to a value of $\sim$1 $\mu$m, consistent with the lower end of the rates of thickness increase observed in experiments. It is possible that increases in thickness increase could occur if the rate of actin polymerization was not constant but accelerated in time—for instance, by depending on the amount of polymerized actin.
- Finally, assuming a typical actin Young's modulus of $E = 0.1$ GPa, actin Poisson's ratio of $\nu = 0$, and an osmotic pressure of $p = 1$ atm (corresponding to an osmolyte concentration gradient of $\sim$50 mM), at long times for which the mechanical strains approximately vanish, the radial shrinkage is predicted to be $\sim$5%.

While the model is quantitatively consistent with the different dynamics observed in the experiments, we note here that it is also possible that differences can arise from the limitations of using a linear theory of elasticity, which assumes small deformations.

#### Predicting eccentricity through mechanical deformations

The above analysis assumed that the vesicle was spherical in order to model actin polymerization dynamics. Our empirical analyses further suggest that there is substantive eccentricity in vesicles (Fig. 3d of the main text). In order to determine whether this eccentricity might arise as a result of asymmetric actin polymerization, we modeled the presence of different actin cap shapes. Fig. 4d of the main text shows the result of Abaqus FEA simulations for strains of the order of $\sim$0.1 and actin caps with different polymerized subtended angles, $\varphi_0$. For these characteristic values, we found considerable eccentricity, on the order of $\sim$0.2 in the deformed shapes.

For all Abaqus FEA analyses performed in this work (as shown in Fig. 4 of the main text), spherical shells of radius $1$, each containing 900 finite elements (a $30 \times 30$ discretization of $\theta, \varphi \in [0, 2\pi] \times [0, \pi])$ were generated and modeled with an elastic material with dimensionless elastic modulus $1$. The baseline thickness of the shell was set to $0.02$, and dimensionless pressure loads of $0.002$ or $0.01$ were applied.

#### Supplementary Figures

##### Supplementary Figure 1

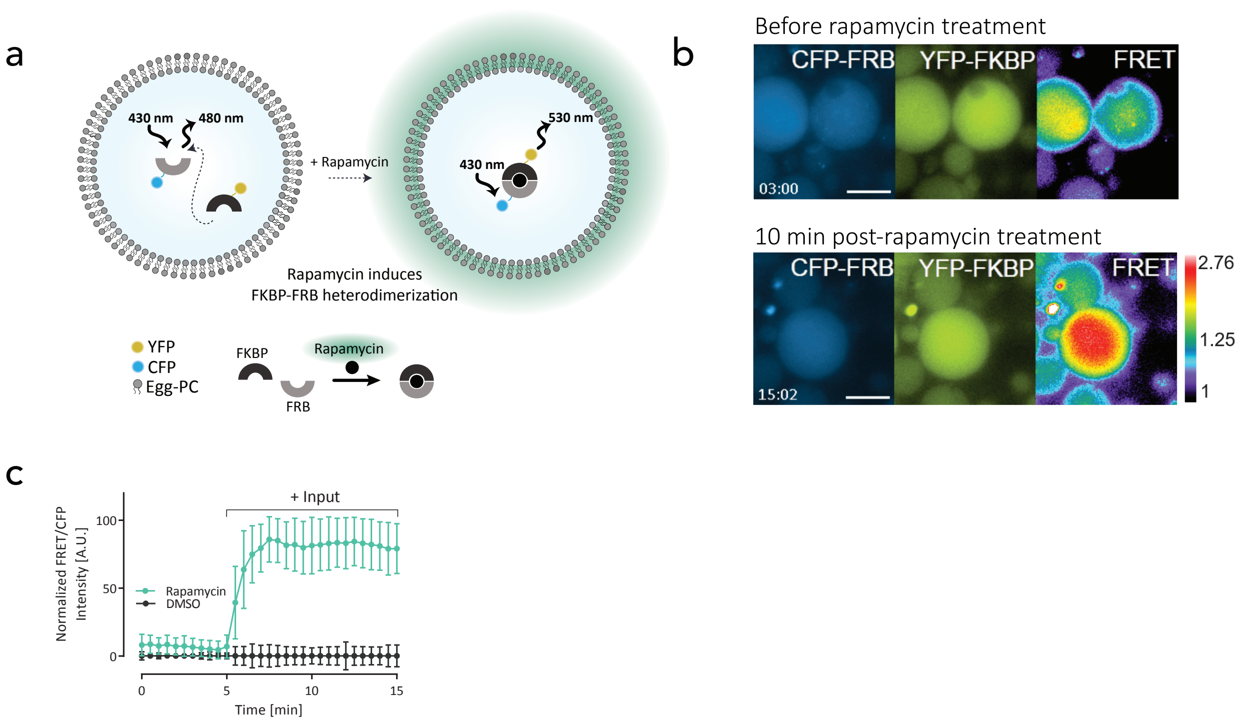


**Supplementary Figure 1.** Reconstitution of chemical sensing inside GUVs. **a.** Schematic of the reconstitution of FKBP and FRB fused with YFP and CFP tags. This protein pair heterodimerizes in the presence of rapamycin resulting in FRET signal increase. **b.** Epi-fluorescence images of purified CFP-FRB [3.2 $\mu M$] and YFP-FKBP [3.2 $\mu M$] proteins diffuse in the lumen of the symmetric egg-PC GUVs which form a complex in the presence of rapamycin as indicated by FRET increase. Rapamycin administration resulted in losing the right non-tethered GUV from the field of view. Scale bar is 20 $\mu m$. **c.** Kinetics of the FRET signal normalized by the donor intensity, highlights FKBP-FRB dimerization in the presence of rapamycin but not the DMSO vehicle. n=35 for rapamycin and n=10 for the DMSO conditions. Error bars represent SDs. Luminal fluorescence intensity of the mCherry-FKBP-MARCKS normalized by the average of the initial values prior to rapamycin treatment shows that due to membrane-anchoring, the localization remains unchanged. n=14 for both rapamycin and DMSO conditions. Error bars represent SDs.

#### Supplementary Figure 2

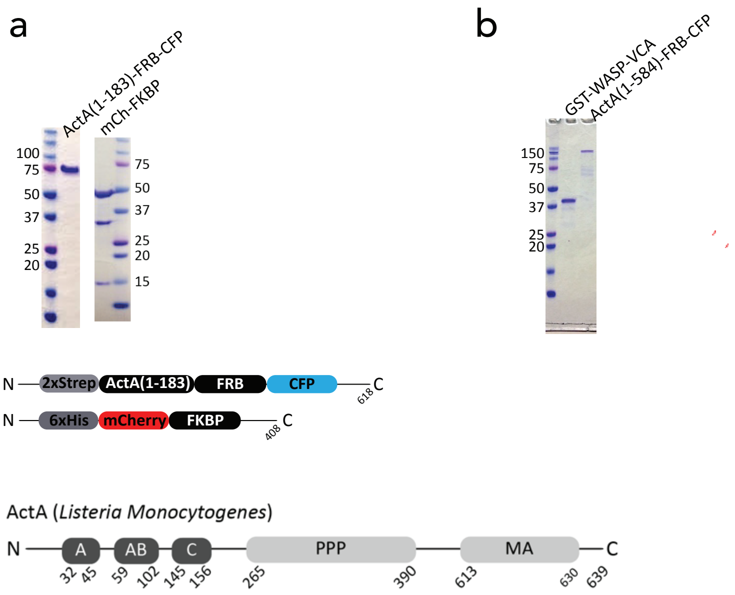


**Supplementary Figure 2.** Protein purification SDS-PAGE gels. **a.** 6xHis-mCh-FKBP and ActA(1-183)-FRB-CFP purified for the CID experiments. **b.** Purified GST-WASP-VCA (purchased from Cytoskeleton) and ActA(1-584)-FRB-CFP used in the pyrene assay.

### Supplementary Figure 3

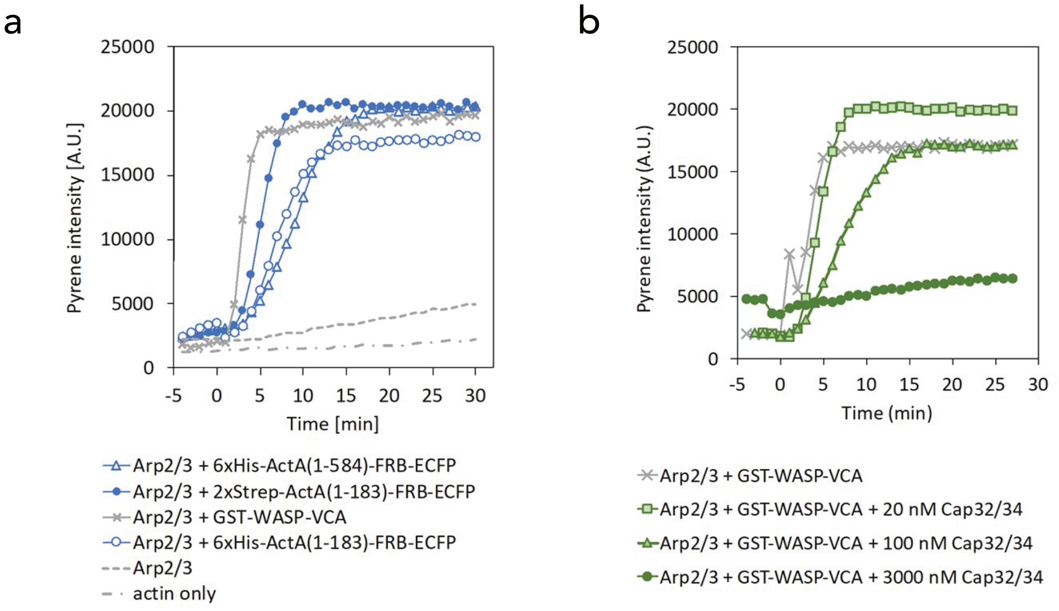


**Supplementary Figure 3.** In vitro actin polymerization activity of ActA. **a.** Pyrene actin assay was carried out to assess actin nucleation promoting factor function of purified ActA. ActA(1-584) and ActA(1-183) and its different purification tag variants were tested for their activity with GST-WASP-VCA as a positive control. The data showed similar activities for all ActA variants. Reaction was started with 1 $\mu M$ actin in F-buffer. Arp2/3 (final 10 $nM$) and GST-WASP-VCA or ActA variants (final 800 $nM$) were added at time = 0. **b.** Activity of purified cap32/34 was evaluated in pyrene actin assay. Cap32/34 reduced actin polymerization rate and the plateau value of F-actin in a dose-dependent manner. Reaction started with 1 $\mu M$ actin with or without cap32/34 in F-buffer and Arp2/3 (final 10 $nM$) and GST-WASP-VCA (final 800 $nM$) were added at time=0.

#### Supplementary Figure 4

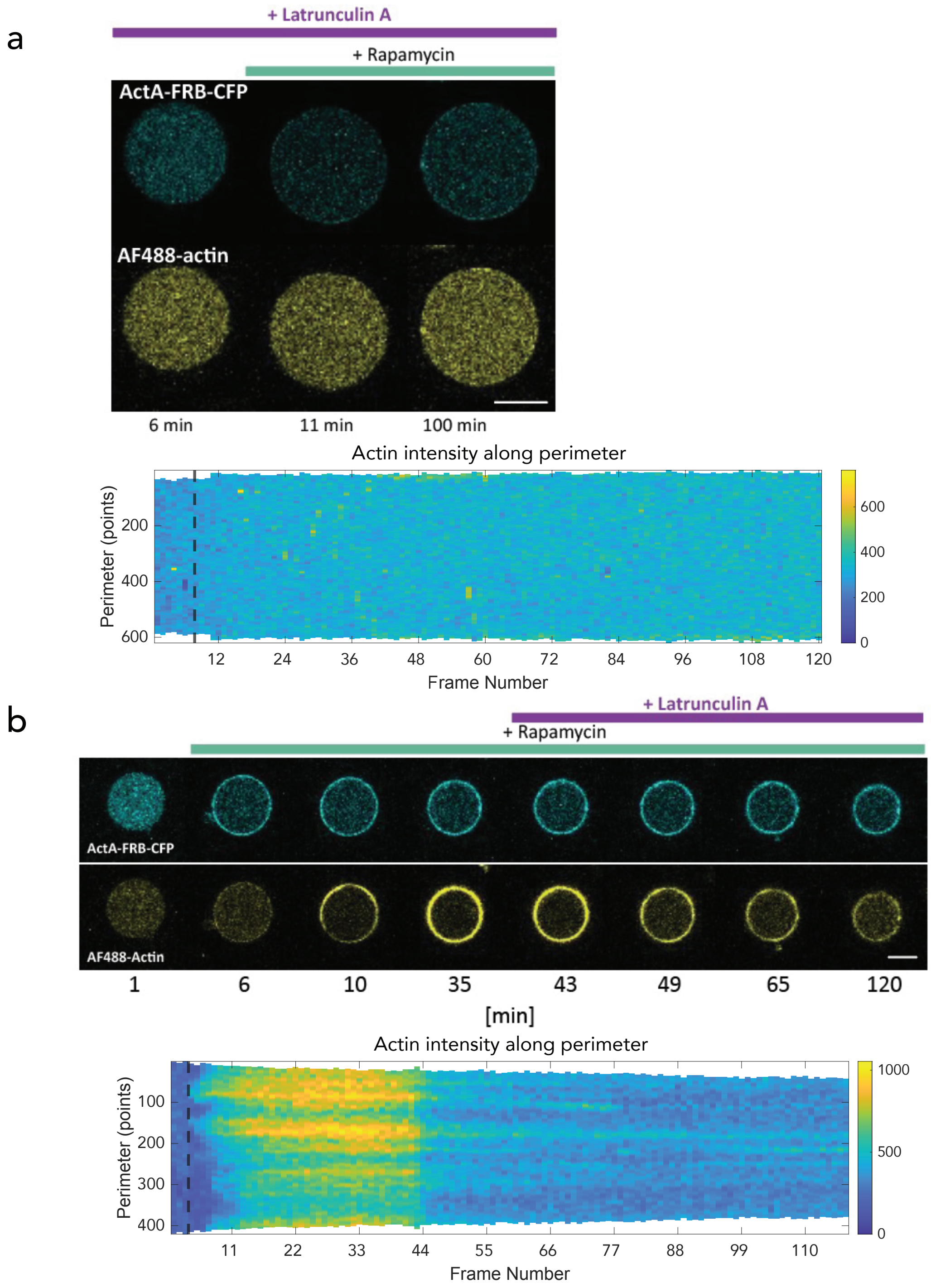


**Supplementary Figure 4.** Latrunculin administration identifies F-actin on the membrane. **a.** With 5 $\mu M$ Latrunculin present, actin does not polymerize on the membrane despite ActA recruitment in the presence of rapamycin. **b.** Alternatively, Latrunculin dissociates the already present F-actin on the membrane.

### Supplementary Figure 5

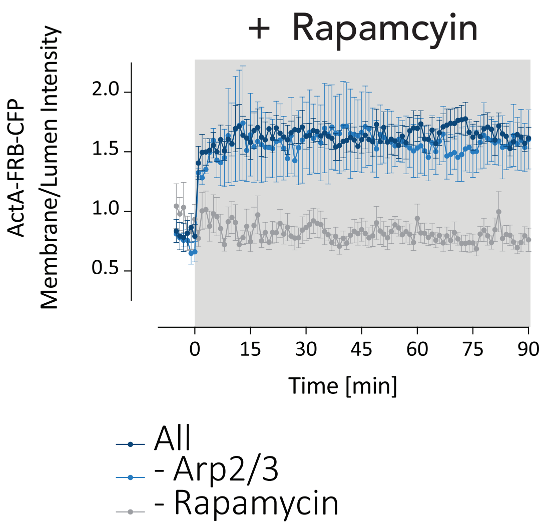


**Supplementary Figure 5.** CID-induced ActA translation towards the membrane. ActA signal increases on the plasma membrane after rapamycin administration in the presence or absence of Arp2/3 complex.

### Supplementary Figure 6

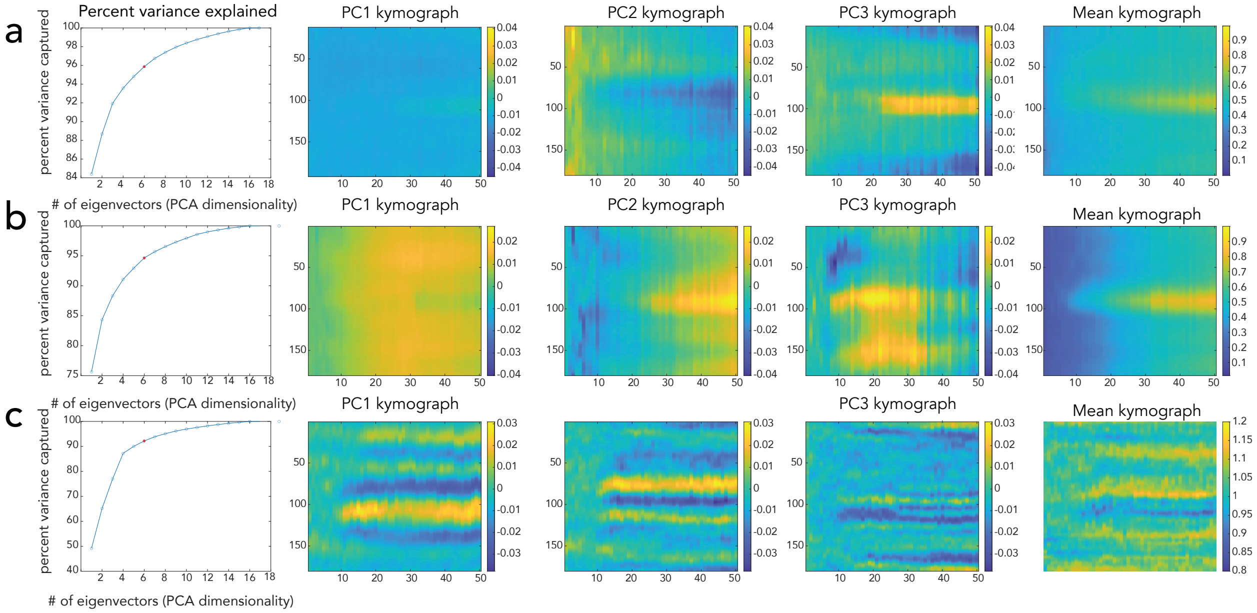


**Supplementary Figure 6.** Principal component analysis (PCA) on the kymographs to extract dominant modes of spatiotemporal dynamics. **a.** PCA performed on ActA kymographs, max-normalized, and aligned based on the actin kymographs at peak actin signal at 35 frames post-rapamycin addition. The top 3 principal components (PCs) capture over 90% of the variance in the dataset. PC1 kymograph represents the dominant spatiotemporal features, followed by PC2, and PC3, relative to the mean kymograph shown. **b.** The same analysis is applied to the actin and **c.** raw curvature kymographs to identify the key spatiotemporal features in actin distribution and local shape changes. n=17 for all panels.

### Supplementary Figure 7

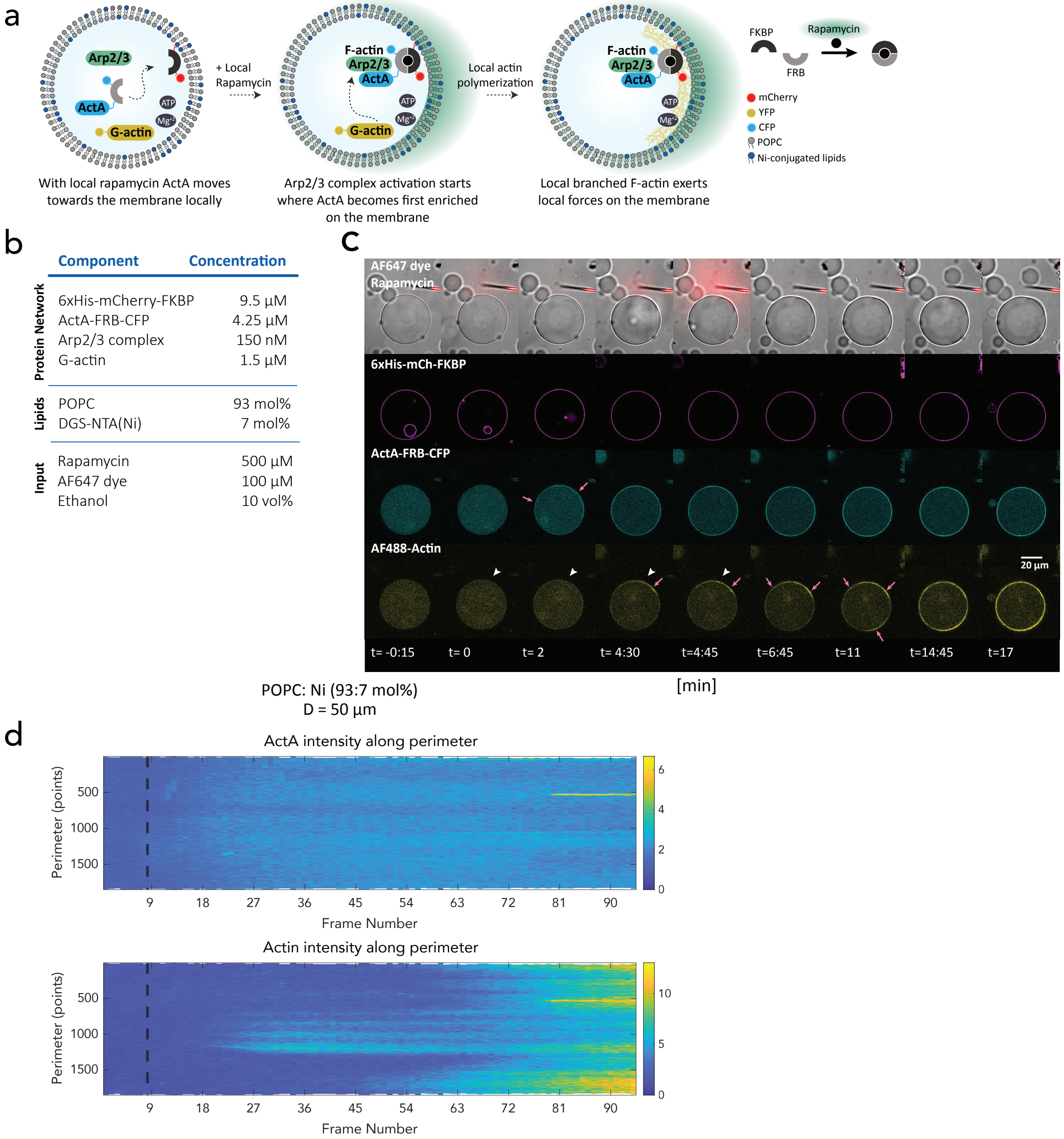


**Supplementary Figure 7.** Local administration of rapamycin for POPC GUVs. **a.** Cartoon depicting a design whereby ActA can translocate locally in response to a rapamycin input, leading to local actin polymerization. **b.** The components assembled in GUVs to mediate local actin polymerization. **c.** With a local rapamycin input, ActA translocated locally and F-actin polymerization initiated next to the rapamycin source, however due to high lipid diffusion ActA and F-actin distribution did not retain their locality. **d.** Kymograph depicting the distribution of ActA and actin on the membrane pre- and post- rapamycin addition.

### Supplementary Figure 8

a

| | Component | Concentration |
|---|---|---|
| Protein Network | 6xHis-mCherry-FKBP | 9.5 µM |
| | ActA-FRB-CFP | 4.25 µM |
| | Arp2/3 complex | 150 nM |
| | G-actin | 1.5 µM |
| Lipids | DPPC | 93 mol% |
| | DGS-NTA(Ni) | 7 mol% |

b

c

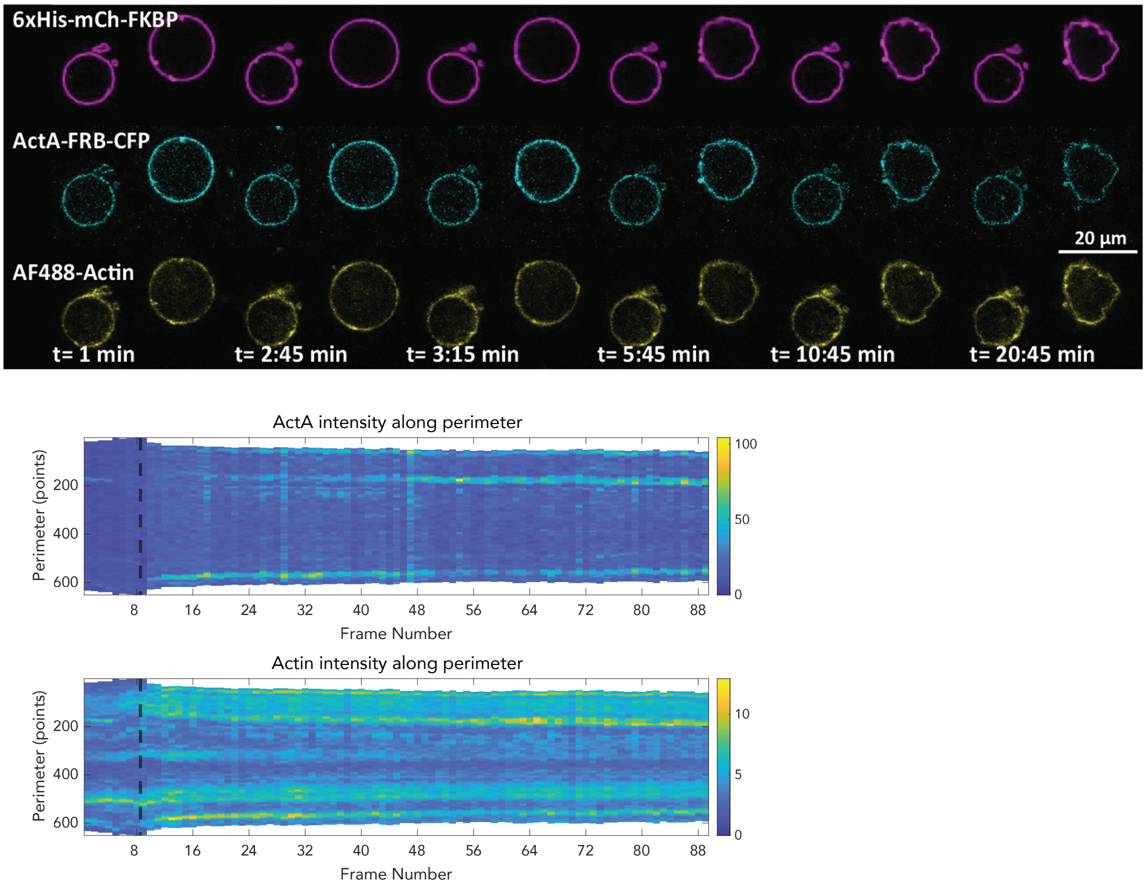


**Supplementary Figure 8.** Global administration of rapamycin for DPPC GUVs. **a.** Assembled GUVs with DPPC lipids and the biomolecules presented. **b.** The more rigid DPPC containing GUVs deformed as a result of actin polymerization on the membrane. **c.** ActA and actin kymographs, depicting the intensity evolution of each on the membrane for the right, larger GUV on (b).

### Supplementary Figure 9

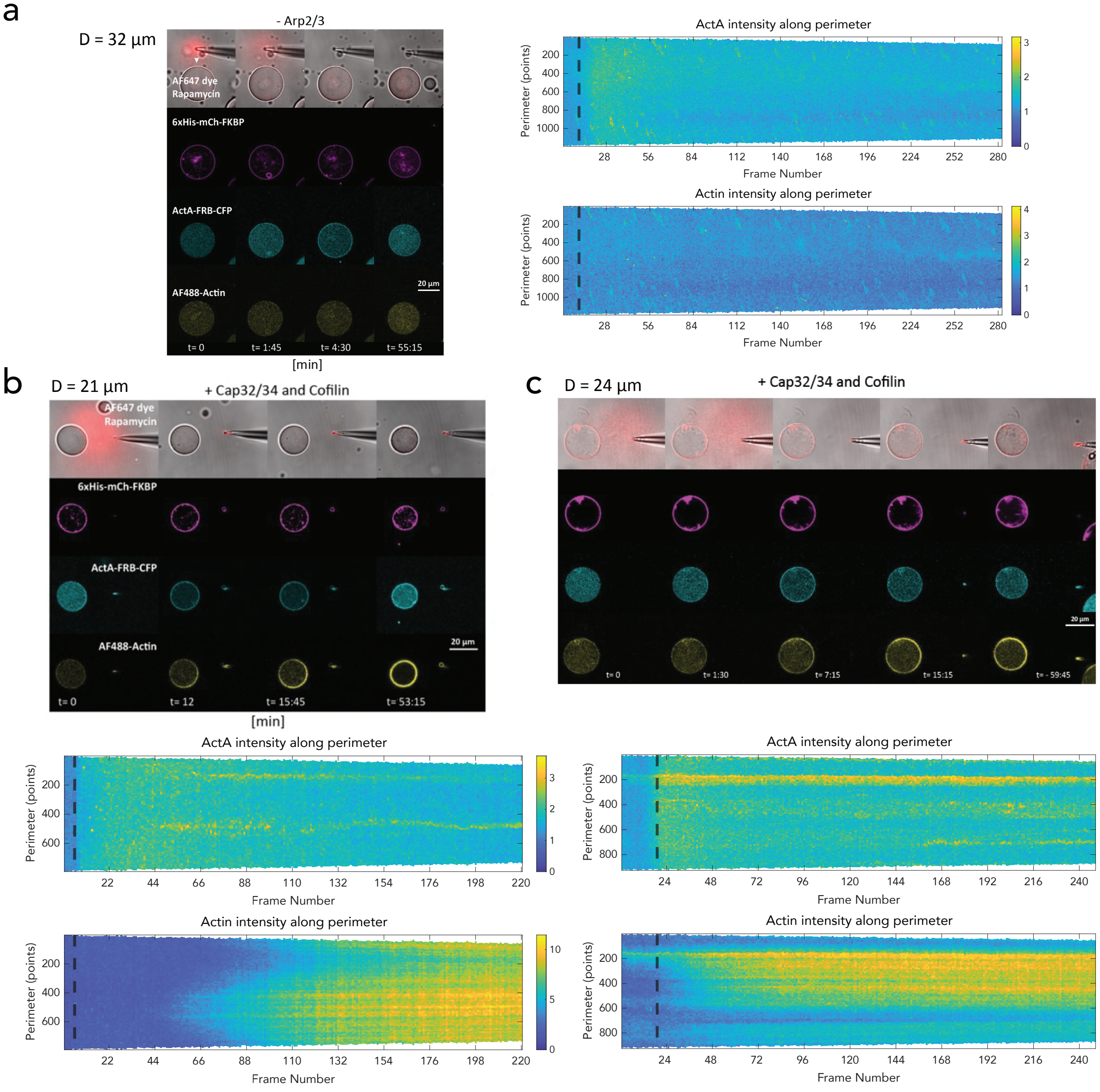


d

| Protein Network | Component | Concentration a | b | c |
|---|---|---|---|---|
| | 6xHis-mCherry-FKBP | 9.5 µM | 9.5 µM | 9.5 µM |
| | ActA-FRB-CFP | 4.25 µM | 4.25 µM | 4.25 µM |
| | Arp2/3 complex | - | 150 nM | 150 nM |
| | G-actin | 1.5 µM | 2 µM | 6 µM |
| | Cap32/34 | - | 50 nM | 50 nM |
| | Cofilin | - | 2 µM | 2 µM |

**Supplementary Figure 9.** Local CID experiments with various biomolecular components. **a.** When no Arp2/3 ActA translocation took place, actin did not polymerize. (**b.-d.**) In the presence of various concentrations of capping protein, cofilin, and actin, the local actin polymerization feature was abolished.

Supplementary Figure 10

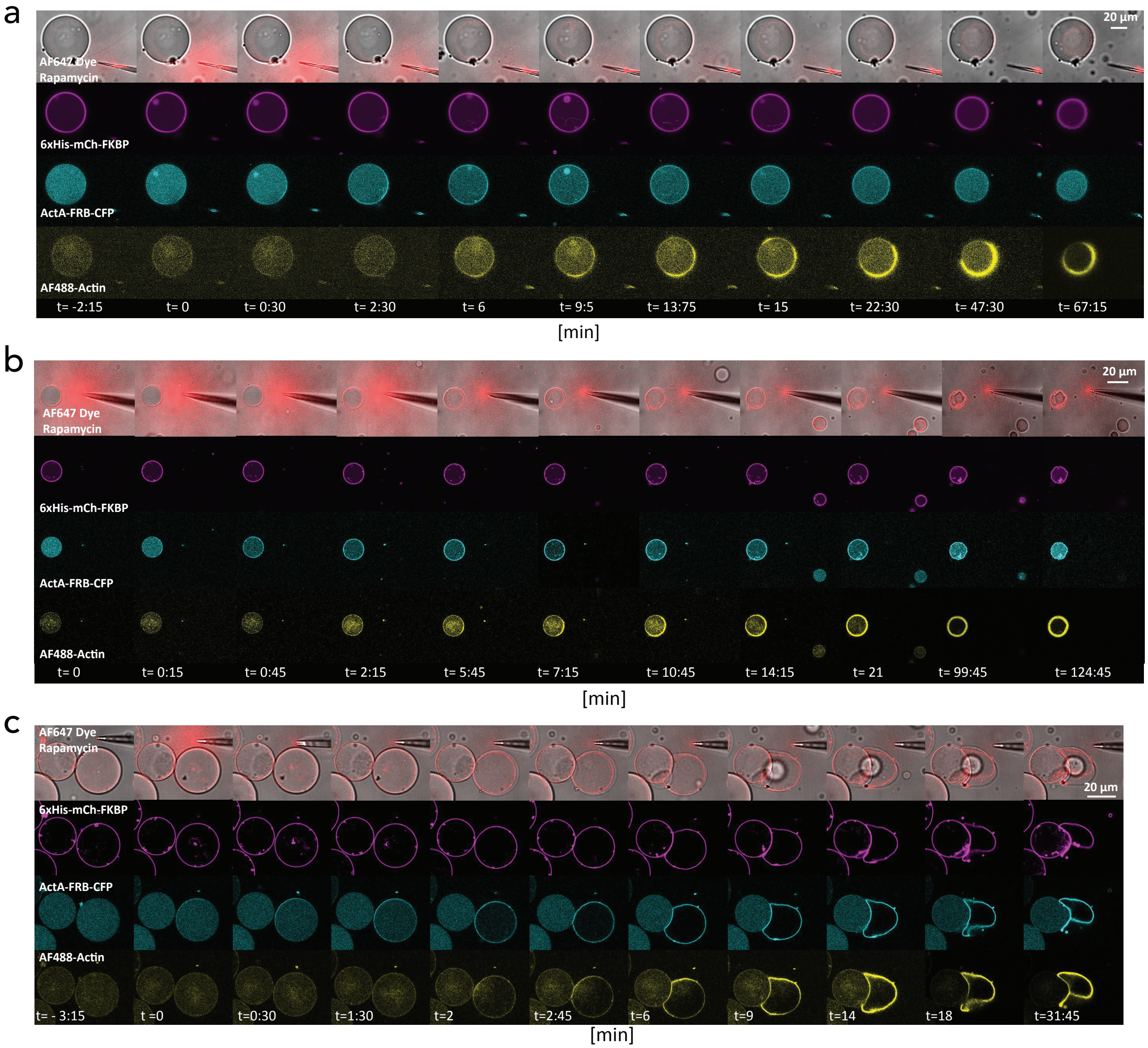


**Supplementary Figure 10.** Representative Local CID experiments. (**a.**-**c.**) Representative images obtained from the local CID experiments (Fig. 5).

### Supplementary Figure 11

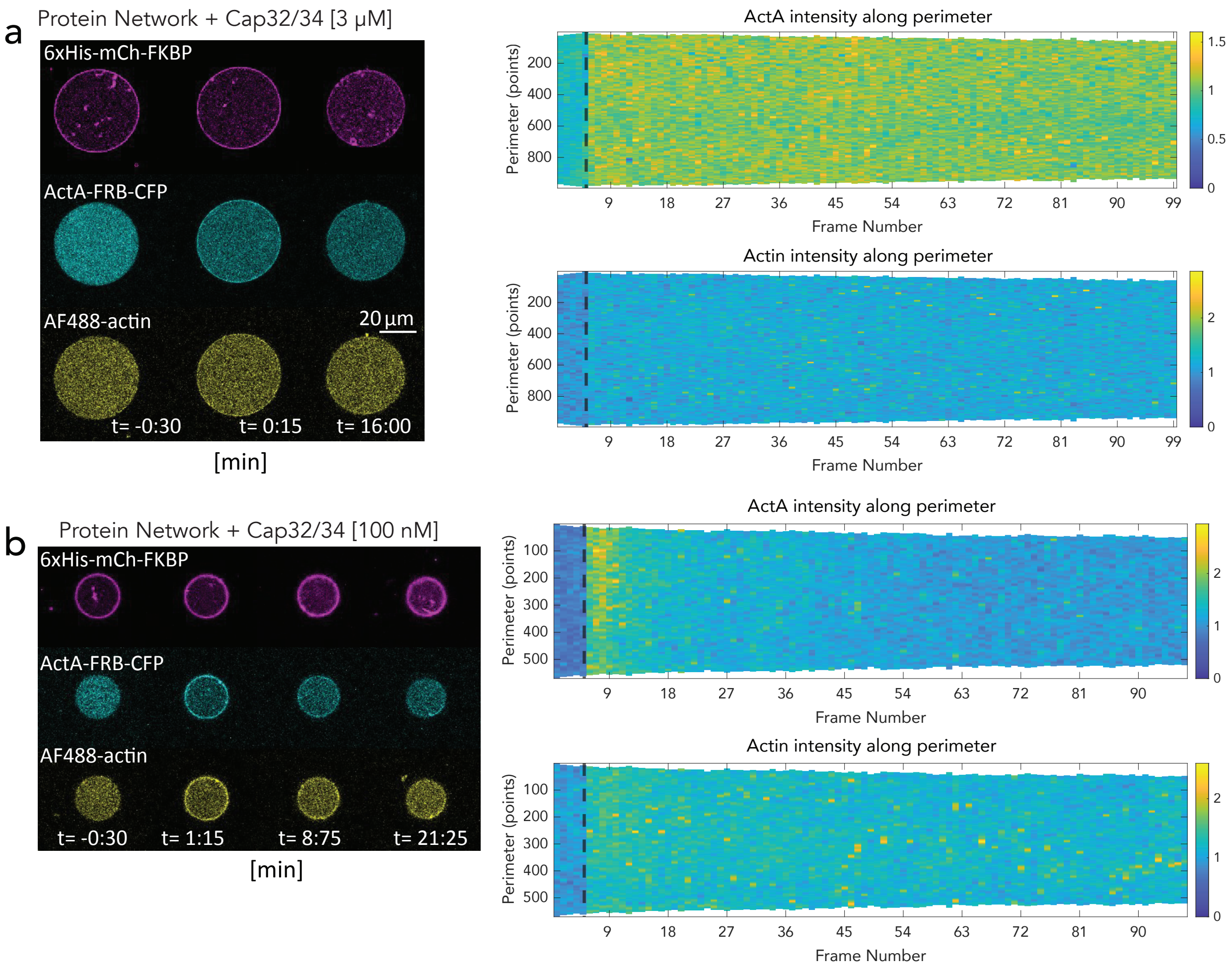


**Supplementary Figure 11.** Cap32/34 and cofilin abolished actin thickening. **a.** Bulk adminstration of rapamycin for GUVs prepared with 3 $\mu M$ Cap32/34. **b.** Bulk administration of rapamycin for GUVs prepared with 100 $nM$ Cap32/34.

#### Supplementary Figure 12

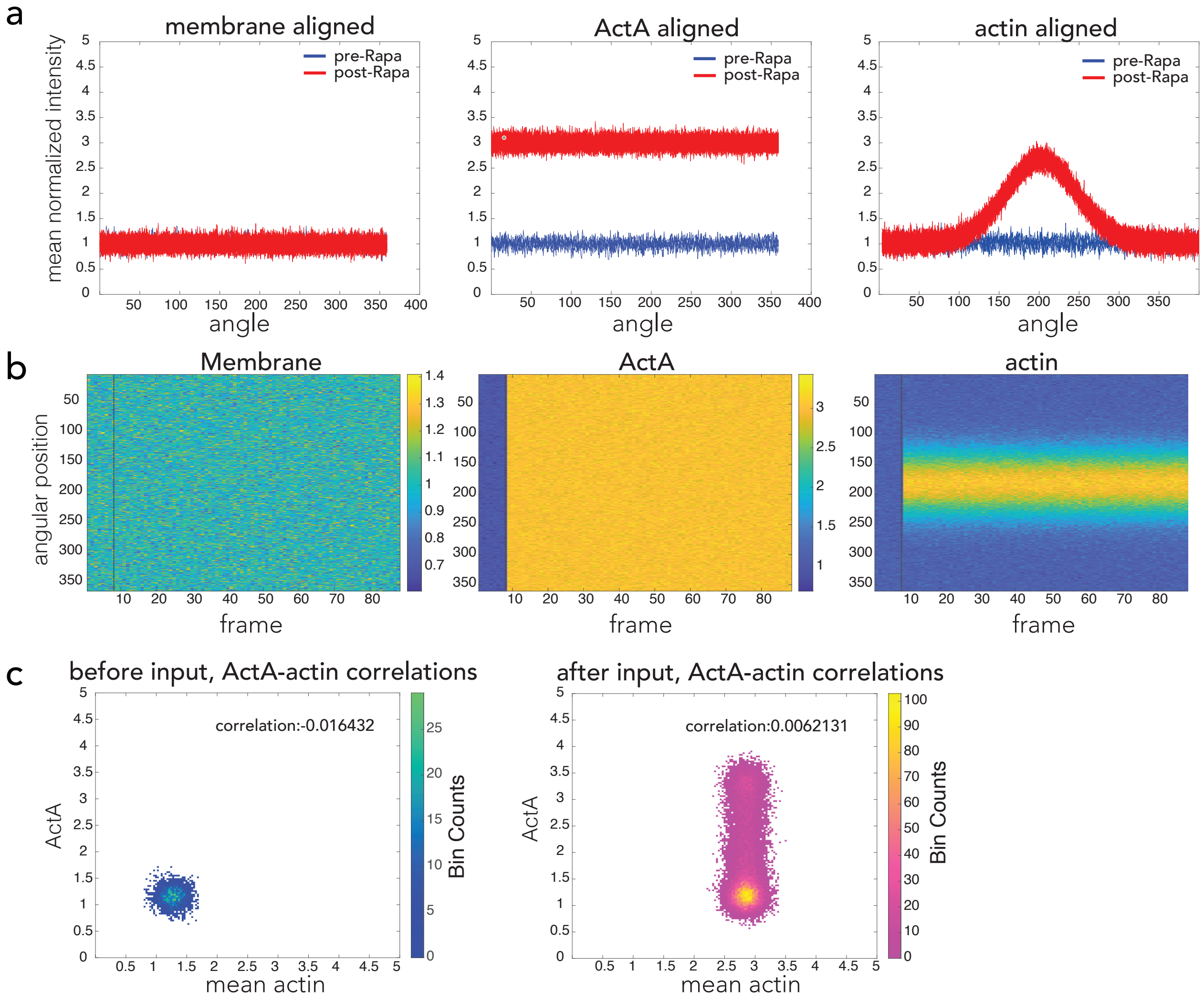


**Supplementary Figure 12.** Assessing our image analysis pipeline against a previously published symmetry breaking report.[43] **a.** Membrane, ActA and actin traces on the GUV boundary before and after actin polymerization on the membrane. **b.** Distribution of membrane, ActA, and actin intensity on the GUV boundary before and after symmetry breaking. **c.** Correlation analysis between ActA and actin before and after actin polymerization on the membrane, highlighting decrease in the Pearson correlation value after actin polymerization induction.

### Supplementary Tables

#### Supplementary Table 1

**Table 1 | Lipid composition of the protocells**

| Figure Number | Leaflet | Lipid | Fatty acid | mol% |
|---|---|---|---|---|
| Fig. 1 | inner | POPC | 16:0-18:1 | 80 |
| | | POPS | 16:0-18:1 | 20 |
| | outer | Egg-PC | N/A | 80 |
| | | POPS | 16:0-18:1 | 20 |
| Figs. 2, 3, Supps. Figs. 4-6, 11 | inner/outer | POPC | 16:0-18:1 | 94 |
| | | DGS-NTA(Ni) | 18:1-18:1 | 5 |
| | | PEG2000 PE | 16:0-16:0 | 1 |
| Fig. 5, Supp. Figs. 8-10 | inner/outer | DPPC | 18:0-18:0 | 93 |
| | | DGS-NTA(Ni) | 18:1-18:1 | 7 |
| Supp. Fig. 1 | inner/outer | Egg-PC | NA | 100 |
| Supp. Fig. 7 | inner/outer | POPC | 16:0-18:1 | 93 |
| | | DGS-NTA(Ni) | 18:1-18:1 | 7 |

POPC: 1-palmitoyl-2-oleyl-sn-glycero-3-phosphocholine
POPS: 1-palmitoyl-2-oleoyl-sn-glycero-3-phospho-L-serine
Egg PC: L-α-phosphatidylcholine (Egg, Chicken)
DGS-NTA(Ni): 1,2-dioleoyl-sn-glycero-3-[(N-(5-amino-1-carboxypentyl)iminodiacetic acid)succinyl]
PEG2000 PE: 1,2-dipalmitoyl-sn-glycero-3-phosphoethanolamine-N-[methoxy(polyethylene glycol)-2000]

**Supplementary Table 1.** Lipid composition of the protocells fabricated for various experimental conditions.

### Supplementary Table 2

##### Global

| | Component | Concentration |
|---|---|---|
| Protein Network | 6xHis-mCherry-FKBP | 9.5 µM |
| | ActA-FRB-CFP | 8.5 µM |
| | Arp2/3 complex | 150 nM |
| | G-actin | 1.5 µM |
| Lipids | POPC | 94 mol% |
| | DGS-NTA(Ni) | 4 mol% |
| | PEG | 1 mol% |

##### Local

| | Component | Concentration |
|---|---|---|
| Protein Network | 6xHis-mCherry-FKBP | 9.5 µM |
| | ActA-FRB-CFP | 4.25 µM |
| | Arp2/3 complex | 150 nM |
| | G-actin | 1.5 µM |
| Lipids | DPPC | 93 mol% |
| | DGS-NTA(Ni) | 7 mol% |
| Input | Rapamycin | 500 µM |
| | AF647 dye | 100 µM |
| | Ethanol | 10 vol% |

**Supplementary Table 2.** Molecular constituents of the protocells used in bulk versus local CID experiments.

#### Supplementary Movies 1 to 3 captions

**Movie S1.** Time-lapse images of the CFP-FRB translocation towards mCh-FKBP-MARCKS upon rapamycin administration.

**Movie S2.** Time-lapse images of rapamycin-induced global ActA translocation towards the membrane and the ensuing actin polymerization that deforms the membrane into an asymmetric shape.

**Movie S3.** Time-lapse images of rapamycin-induced localized ActA translocation towards the membrane and the ensuing local actin polymerization.
